# Single Molecule Analysis of CENP-A Chromatin by High-Speed Atomic Force Microscopy

**DOI:** 10.1101/2022.01.04.474986

**Authors:** Daniël P. Melters, Keir C. Neuman, Tatini Rakshit, Yamini Dalal

## Abstract

Chromatin accessibility is modulated in a variety of ways to create open and closed chromatin states, both of which are critical for eukaryotic gene regulation. At the single molecule level, how accessibility is regulated in the chromatin fiber composed of canonical or variant nucleosomes is a fundamental question in the field. Here, we developed a single-molecule tracking method where we could analyze thousands of canonical H3 and centromeric variant nucleosomes imaged by high-speed atomic force microscopy. This approach allowed us to investigate how changes in nucleosome dynamics *in vitro* inform us about chromatin accessibility *in vivo*. By high-speed atomic force microscopy, we tracked chromatin dynamics in real time and determined the MSD and diffusion constant for the variant centromeric CENP-A nucleosome. Furthermore, an essential kinetochore protein CENP-C reduces the diffusion constant and mobility of centromeric nucleosomes along the chromatin fiber. We subsequently interrogated how CENP-C modulates CENP-A chromatin dynamics *in vivo*. Overexpressing CENP-C resulted in reduced centromeric transcription and impaired loading of new CENP-A molecules. Thus, changes which alter chromatin accessibility *in vitro*, also correspondingly alter transcription *in vivo*. These data suggest a model in which variant nucleosomes encode their own diffusion kinetics and mobility, and where binding partners can suppress or enhance mobility.

## Introduction

Regulating physical access to DNA is central to both gene expression, genome topology, and genome integrity across eukaryotes. Decades of data strongly suggest that more physically accessible chromatin is also more transcriptionally permissive (Klemm et al., 2019; Maeshima et al., 2019; and landmark papers from Weintraub and Groudine, 1976; Wu et al., 1979). In contrast, compacted chromatin or heterochromatin has been correlated with transcriptional restriction (Allshire and Madhani, 2018; Flamm et al., 1969; Janssen et al., 2018; Schultz, 1936). These chromatin states are dynamic and subject to tight regulation. Chromatin-binding proteins dictate many of these dynamics in part driven by the presence and deposition of specific post-translational modifications (PTMs) of nucleosomes (Chung et al., 2023; Rothbart and Strahl, 2014; Taverna et al., 2007; Tolsma and Hansen, 2019). For instance, HP1 binds to H3K9me2/3 nucleosomes (Bannister et al., 2001; Jun-ichi et al., 2001; Lachner et al., 2001; Sanulli et al., 2019a), ultimately resulting in transcriptionally repressive chromatin (Bannister et al., 2001; Escobar et al., 2021; Hwang et al., 2001; Jun-ichi et al., 2001; Lachner et al., 2001).

Interestingly, upon binding of HP1 to H3K9me3 nucleosomes, these nucleosomes’ internal residues become more exposed to hydrogen/deuterium exchange (Sanulli et al., 2019b). This seminal finding suggests that chromatin-binding proteins not only serve as a recruitment platform for other binding proteins but can rapidly alter the physical properties of individual nucleosomes. The logical extension of this concept is examining how altering innate physical properties of nucleosomes modulates the local chromatin state’s structure and function.

To study how nucleosome dynamics is altered by chromatin binding factors, single molecule techniques have been developed, ranging from groundbreaking *in vitro* techniques such as optical and magnetic tweezers (Bustamante et al., 2021; Chien and van Noort, 2009; Killian et al., 2018; Neuman and Nagy, 2008) and *in vivo* single molecule tracking (Iida et al., n.d.; Izeddin et al., 2014; Mueller et al., 2013; Shen et al., 2017; Wang et al., 2021). The former two techniques are exclusively *in vitro* and rely on precisely designed DNA sequences and constructs to guarantee precise measurements. By manipulating an optical trap or magnetic tweezer, tension and torsion forces can be exerted on the associated DNA molecule, which in turn alters the forces exerted on nucleosomes (Bustamante et al., 2021; Chien and van Noort, 2009; Killian et al., 2018; Neuman and Nagy, 2008). In contrast, single molecule tracking in cells is made possible by photostable fluorophores covalently bound to a target protein. These tagged proteins are introduced into cells at low concentration to allow the tracking of single molecules, with limited control where the tagged proteins will go (Iida et al., n.d.; Mueller et al., 2013; Shen et al., 2017; Wang et al., 2021). Both systems are powerful and have distinct advantages, ranging from bp-precision of nucleosome sliding or folding-unfolding dynamics to determining residency time of transcription factors. However, a gap exists connecting these two technical approaches, namely assessing, and quantifying the dynamics of individual nucleosomes and correlating them with global chromatin dynamics. High-speed atomic force microscopy (HS-AFM) has the capability to cover this gap. It is an *in vitro* based technique that permits real time tracking of single molecules in the context of interacting macromolecular complexes (e.g. myosin tracking on actin filaments, Ando, 2018; chromatin techniques reviewed in Melters and Dalal, 2021). By imaging nucleosome arrays in buffer over time, it is possible not just to track, but also quantify the motions of individual nucleosomes within an array.

In addition to PTMs, the chromatin landscape is also marked by the local enrichment of histone variants (Buschbeck and Hake, 2017; Martire and Banaszynski, 2020; Melters et al., 2015), such as the centromere-specific H3 histone variant CENP-A/CENH3. CENP-A nucleosomes recruit several centromeric proteins (Kai et al., 2021; Mendiburo et al., 2011; Régnier et al., 2005) including CENP-C. CENP-C in turn functions as the blueprint for the formation of the kinetochore (Cheeseman et al., 2006; DeLuca and Musacchio, 2012; Kai et al., 2021; Przewloka et al., 2007; Weir et al., 2016). Recently, we reported that a CENP-C fragment (CENP-C^CD^) induces loss of CENP-A nucleosomal elasticity *in silico* and *in vitro* (Melters et al., 2019). This finding correlates with decreased hydrogen/deuterium exchange of CENP-A nucleosomes when bound by the central domain of CENP-C (CENP-C^CD^; Falk et al., 2016, 2015; Guo et al., 2017). Interestingly, when CENP-C was overexpressed, RNA polymerase 2 (RNAP2) levels at the centromere were reduced and centromeric chromatin clustered together (Melters et al., 2019).

We were curious about how CENP-A alone, and in combination with CENP-C mechanistically impacts centromeric chromatin fiber mobility and accessibility in real time. To address this question, we employed HS-AFM to track thousands of canonical or centromeric nucleosomes in arrays over time *in vitro*. We report that HS-AFM imaging is free of tip-induced artifacts and CENP-A chromatin responds predictably to various control conditions. Next, we find that the essential kinetochore protein CENP-C, which is CENP-A chromatin’s closest binding partner, directly impacts nucleosome mobility and, surprisingly, also chromatin fiber motion *in vitro*.

These data represent a technological advance in imaging and analyzing chromatin dynamics by HS-AFM. We extended these findings *in vivo* using immunofluorescence imaging and biochemical approaches, reporting that over-expressing CENP-C alters centromeric chromatin transcription and the ability to load new CENP-A molecules. Cumulatively, these data provide the first direct mechanistic evidence for local transcriptional competency dependent on innate properties and local homeostasis of histone variants within the chromatin fiber *in vivo*.

## Results

We are interested in understanding how nucleosomes “behave” in biologically relevant conditions. Single-molecule techniques have enabled us to understand details about the movement of transcription factors inside the nucleus (Iida et al., n.d.; Mueller et al., 2013; Shen et al., 2017; Wang et al., 2021) and how torsion and pulling/pushing forces influence nucleosomes (Bustamante et al., 2021; Chien and van Noort, 2009; Killian et al., 2018; Neuman and Nagy, 2008). The behavior of a single trajectory might be stochastic, the statistical behavior from many trajectories may reveal additional physical properties, such as diffusion and folding-unfolding dynamics.

By directly observing topographic characteristics and dynamics of chromatin using HS-AFM, we were able to assess the motions of individual nucleosomes in real-time (Figure 1). This emerging single-molecule technique is powerful, and shares similarities with live cell imaging (Ashwin et al., 2019; Specht et al., 2017), magnetic tweezers, and optical tweezers (Bustamante et al., 2021; Chien and van Noort, 2009; Killian et al., 2018; Neuman and Nagy, 2008). Whereas, live cell imaging and single-molecule force spectroscopy methods rely on fluorophore-tags and tethering, HS-AFM can be done on both unmodified and modified protein all while requiring minimal sample preparation (Ando, 2018).

**Figure 1.**
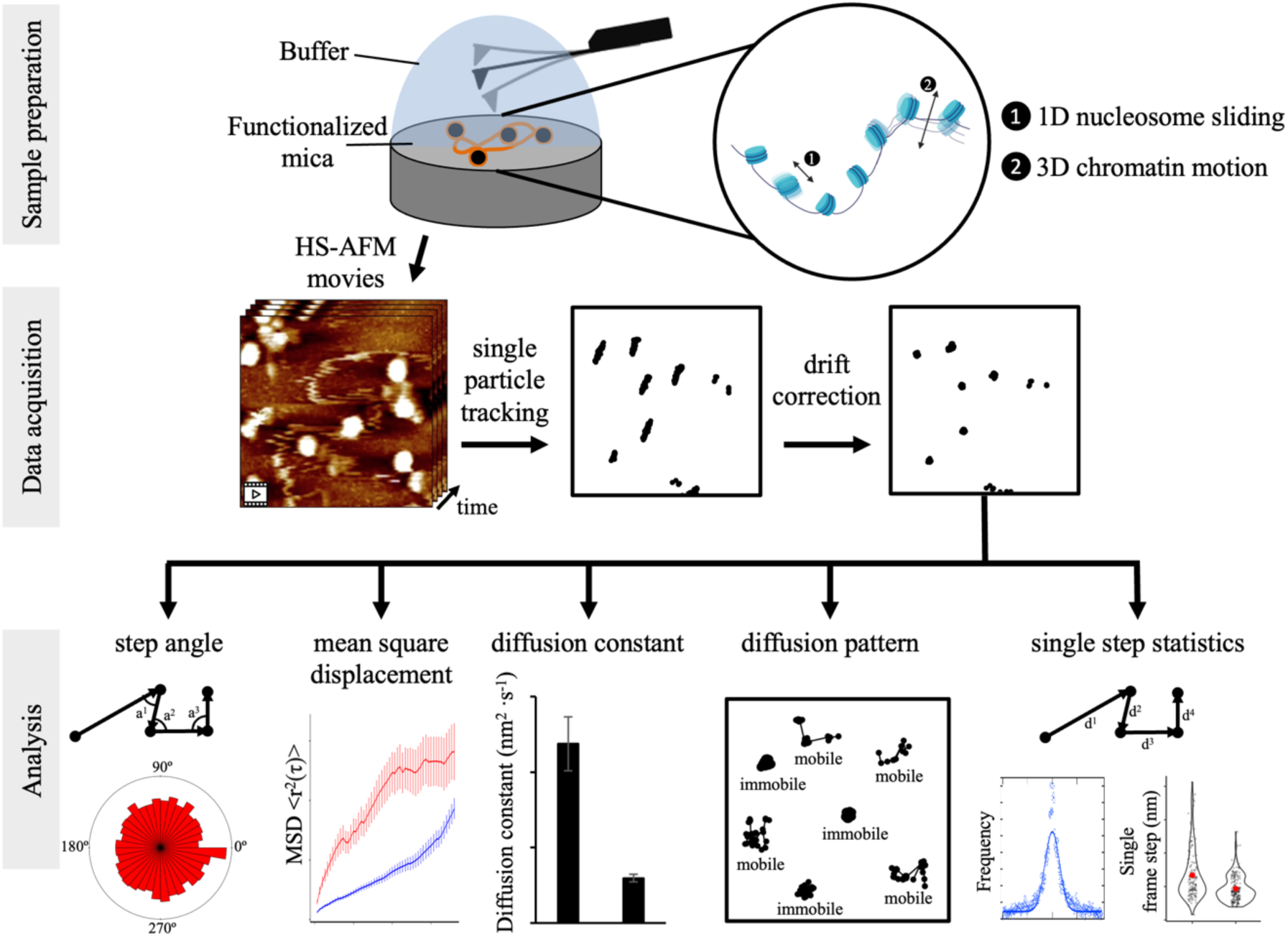
Schematic of experimental set-up for HS-AFM nucleosome measurements and analysis of extracted single particle trajectories. Sample preparation: CENP-A nucleosomes were *in vitro* reconstituted and imaged by HS-AFM in fluid. By HS-AFM we can track nucleosome motion, corresponding to both nucleosomes sliding along the DNA and chromatin fiber moving as a unit. Data acquisition: HS-AFM movies were obtained at a framerate of 0.5 Hz (2 s per frame) for a minimum of 20 seconds and up to 120 seconds with a maximum single-frame step of 24 nm per particle trajectory. Using MATLAB we extracted nucleosome trajectories, which were subsequently corrected for drift. Analysis: trajectories were analyzed to extract several parameters to determine both potential tip-scanning artifacts and to characterize nucleosome dynamics.

Recently, we showed by HS-AFM that, at a qualitative level, H3 nucleosomes are mobile and make intermittent contact with other H3 nucleosomes (Melters and Dalal, 2021). Here, we set out to quantify the potential dynamics of individual nucleosomes in the context of chromatin. This means that we observe and quantify global nucleosome movement, which can be 1- dimenstional motion of nucleosome sliding along DNA events as well as 3-dimensional whole chromatin fiber movements (Figure 1). We predict that these global nucleosome motions reflect the complex dynamics nucleosomes display in the nucleus (Ide et al., 2022; Iida et al., n.d.). To assay CENP-A chromatin, we reconstituted CENP-A nucleosomes on a ∼3.5 kbp plasmid containing four copies of *α*-satellite repeats. After verifying successful *in vitro* chromatin reconstitution (Supplemental Figure 1, Source data 1), we imaged CENP-A chromatin using the Cypher VRS system (see material and methods for details). Movies were obtained at two seconds per frame. Next, the movies were converted to TIFF sequences. Using MATLAB’s single particle tracking package, we obtained a total of 4,374 individual nucleosome trajectories from 10 different samples (see Material and Methods for details). To obtain useful mean square displacement (MSD) values, individual trajectories needed to be of sufficient duration.

Therefore, we selected individual trajectories that were at least 20 seconds, with a maximum of 120 seconds (10-60 points). In addition, we wanted to make sure that we were tracking the same nucleosomes, so we estimated the maximum single step size at twice the nucleosome width or 24 nm. Next, we corrected each movie for drift (Supplemental Figure 2). Finally, we analyzed 2,591 trajectories to obtain the step angle, step size, MSD, and diffusion constant of individual nucleosomes (Figure 1).

### HS-AFM imaging is free of tip-induced artifacts

AFM is a topographical imaging technique, and a common concern is that the AFM tip may alter the sample during scanning. To determine whether there is indeed a tip effect, we performed several controls. We reasoned that if the AFM tip altered samples during scanning, it would create a distinctive signature in single particle trajectories. We imaged CENP-A chromatin under multiple conditions (Supplemental movies 1-5). First, we assessed the angle between successive steps (Figure 2A). If there was a tip effect or drift, there would be a bias in the distribution of angles. We did not observe a bias in the angle distributions (Figure 2B; Supplemental Figure 3A). Second, we looked at the diffusion constant over the x-axis alone or the y-axis alone (Figure 2C) in the absence or presence of 1-(3-aminopropyl) silane (APS). APS functionalizes the mica surface with positively charged amino groups, binding nucleic acid molecules under physiological conditions, allowing for the ability to image in air, in fluid, as well as perform force spectroscopy (Lyubchenko et al., 2014; McAllister et al., 2005; Melters et al., 2019; Melters and Dalal, 2021; Shlyakhtenko et al., 2003). We predicted that adding an excess of APS (333 nM APS is 2x APS) would lower the diffusion constant compared to not adding APS (no APS). Indeed, we detected a lower diffusion constant for 2x APS than no APS (Figure 2D, Source data 2, 3). If there were a tip effect, we reasoned there would be a difference in the diffusion constants between the x-axis and y-axis, which are parallel and perpendicular to the direction of tip scanning, respectively. We did not observe a bias in the diffusion constants between the two axes (Figure 2D; Supplemental Figure 3B, Source data 2, 3). Third, to test for the potential impact of prolonged imaging on manipulation of trajectories, we assessed the step size distribution over time. We did not observe a significant effect of the movie length on the step size (Supplemental Figure 4, Source data 2), indicating that extended imaging did not alter the trajectory dynamics. We are therefore confident that HS-AFM does not introduce a tip effect on chromatin and that drift is adequately corrected.

**Figure 2.**
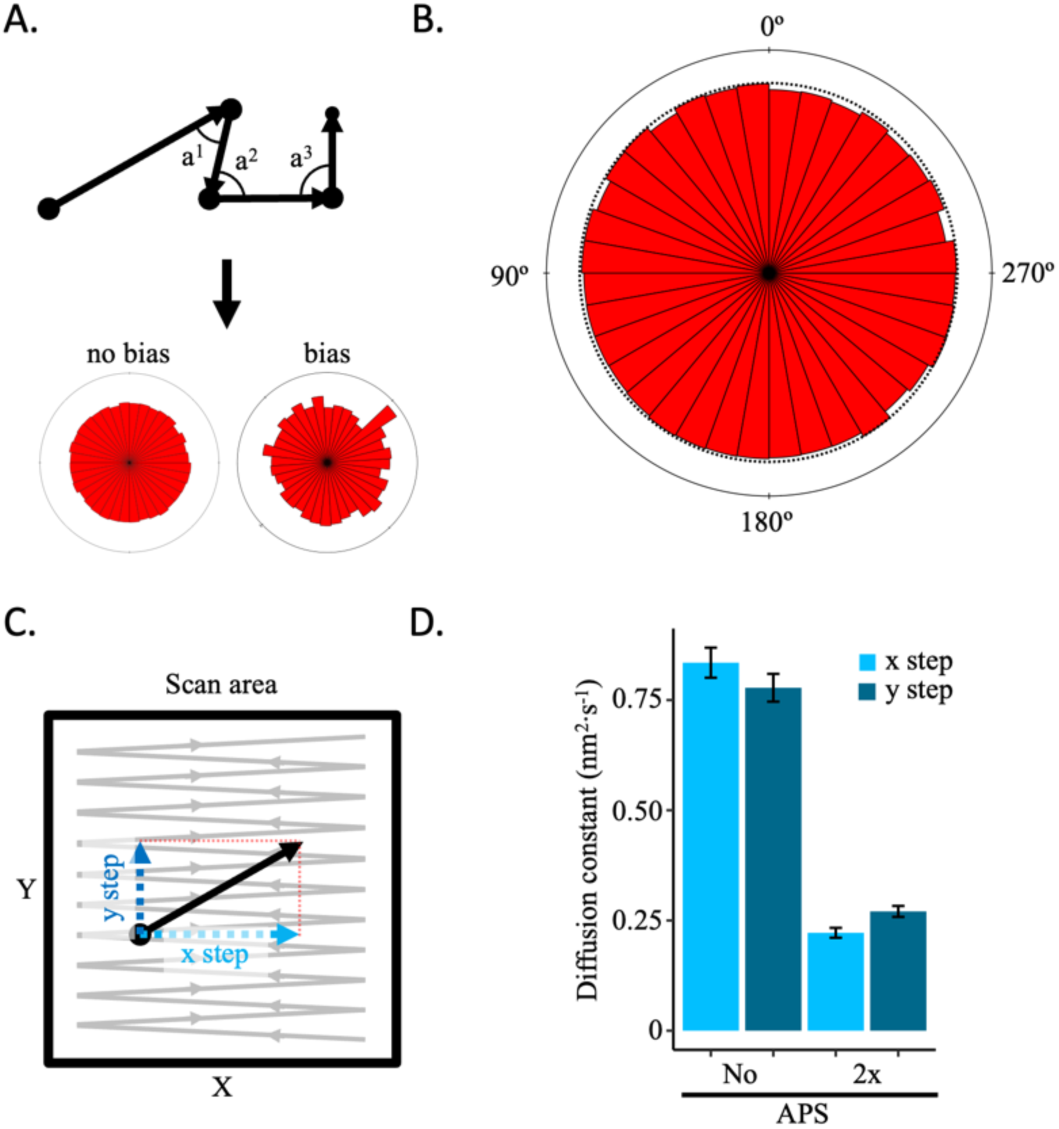
No AFM tip effect was observed. If the AFM tip were to displace the sample during scanning, it should result in a motion bias in the direction of scanning that can be detected. (A) Schematic representation of how angle between successive steps within a trajectory is determined and representative angle distribution graphs for no bias or bias. (B) All angles for successive steps of all trajectories of five control conditions (low salt, high salt, no APS, 2x APS, and Tween-20) show no sign of bias (Supplementary Figure 2A). (C) Every step has an x and y coordinate. By obtaining the diffusion constant for each axis separately, motion bias between the x and y axis can be discerned, which is an indication for bias introduced by the AMF tip. (D) The diffusion constants for the x and y axis for CENP-A nucleosomes in low or high salt conditions shows a difference between imaging conditions, but not within each condition between the x and y axis (Supplemental Figure 2B).

### Salt and APS concentrations impact chromatin dynamics in an anticipated manner

Next, we set out to determine how different buffer conditions impact CENP-A chromatin dynamics. Different salt concentrations are known to impact chromatin compaction and dynamics (Allahverdi et al., 2015; Brasch et al., 1971; Yager et al., 1989; Yager and Van Holde, 1984). At lower salt concentrations (below 50 mM NaCl), nucleosomes stabilize, whereas at higher salt concentrations (above 100 mM NaCl) nucleosomes become more unstable. Here, we tested the effect of low salt concentrations (5mM NaCl) versus high salt concentration (150 mM NaCl) on nucleosome dynamics by HS-AFM. For each condition we tracked 124 and 161 nucleosome trajectories, respectively (Table 1, Supplemental Movie 1, 2, Source data 2). As expected, the MSD curve for high salt had a larger slope than for low salt (Figure 3A, Supplemental Figure 5A, B). This is reflected in the diffusion constant, which was 1.2±0.2 nm^2^·s^-1^ for low salt and 4.1±0.3 nm^2^·s^-1^ for high salt (Figure 3B, Supplemental Figure 6C).

**Figure 3.**
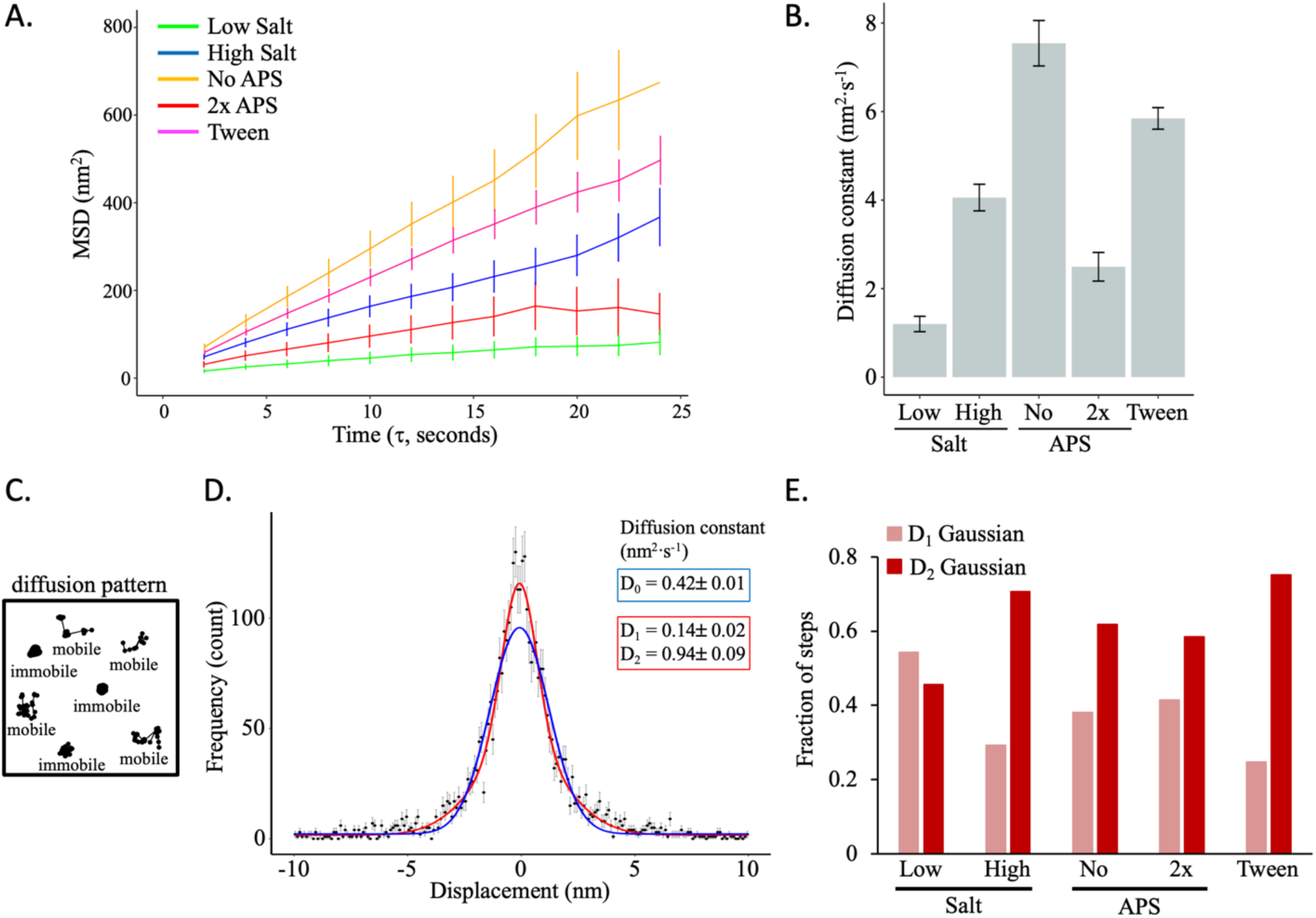
Salt and APS concentration predictably impacts CENP-A nucleosome mobility *in vitro*. (A) The average mean square displacement is shown with standard error as a function of the time interval for CENP- A nucleosome arrays in the following buffers: low salt (green; 5 mM NaCl), high salt (blue, 150 mM NaCl), no APS (yellow), 2-fold APS (red), and 0.01% Tween-20 (pink). (B) The diffusion constants obtained from the MSD curves. (C) Schematic representation of mobile or immobile (or paused) single particle trajectories. (D) The single step x- axis displacement of CENP-A nucleosomes in low salt conditions. The blue line represent a single Gaussian fit and the red line represents a double Gaussian fit. The latter provided a better fit to all the data for both the x and y for all conditions (see Supplemental Figure 5). (E) The fraction of single steps corresponding to D1 (narrower) Gaussian distribution or D2 (wider) Gaussian distributions from the double Gaussian fitting. The D1 Gaussian distribution corresponds to a smaller diffusion constant and may represent immobile or paused nucleosomes, whereas D2 corresponds to a larger diffusion constant representing mobile nucleosomes. The data were obtained from two independent technical replicates per condition.

**Table 1.**
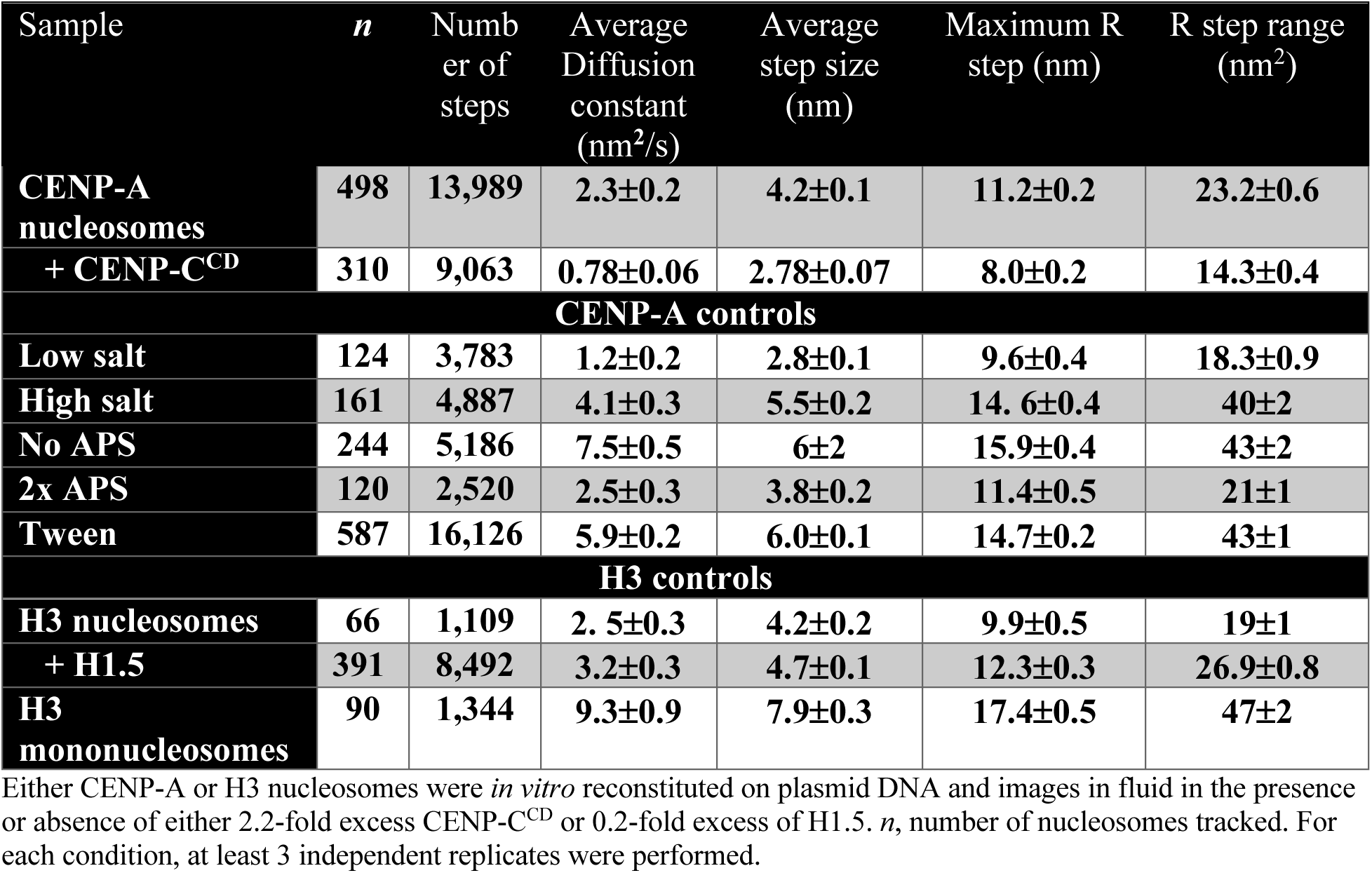
Quantifications of HS-AFM movies

| Sample | *n* | Number of steps | Average Diffusion constant (nm<sup>2</sup>/s) | Average step size (nm) | Maximum R step (nm) | R step range (nm<sup>2</sup>) |
| --- | --- | --- | --- | --- | --- | --- |
| **CENP-A nucleosomes** | **498** | **13,989** | **2.3±0.2** | **4.2±0.1** | **11.2±0.2** | **23.2±0.6** |
| **+ CENP-C<sup>CD</sup>** | **310** | **9,063** | **0.78±0.06** | **2.78±0.07** | **8.0±0.2** | **14.3±0.4** |
| **CENP-A controls** |  |  |  |  |  |  |
| **Low salt** | **124** | **3,783** | **1.2±0.2** | **2.8±0.1** | **9.6±0.4** | **18.3±0.9** |
| **High salt** | **161** | **4,887** | **4.1±0.3** | **5.5±0.2** | **14.6±0.4** | **40±2** |
| **No APS** | **244** | **5,186** | **7.5±0.5** | **6±2** | **15.9±0.4** | **43±2** |
| **2x APS** | **120** | **2,520** | **2.5±0.3** | **3.8±0.2** | **11.4±0.5** | **21±1** |
| **Tween** | **587** | **16,126** | **5.9±0.2** | **6.0±0.1** | **14.7±0.2** | **43±1** |
| **H3 controls** |  |  |  |  |  |  |
| **H3 nucleosomes** | **66** | **1,109** | **2.5±0.3** | **4.2±0.2** | **9.9±0.5** | **19±1** |
| **+ H1.5** | **391** | **8,492** | **3.2±0.3** | **4.7±0.1** | **12.3±0.3** | **26.9±0.8** |
| **H3 mononucleosomes** | **90** | **1,344** | **9.3±0.9** | **7.9±0.3** | **17.4±0.5** | **47±2** |

Either CENP-A or H3 nucleosomes were *in vitro* reconstituted on plasmid DNA and images in fluid in the presence or absence of either 2.2-fold excess CENP-C<sup>CD</sup> or 0.2-fold excess of H1.5. *n*, number of nucleosomes tracked. For each condition, at least 3 independent replicates were performed.

To better understand the step dynamics, we first assessed the average single-frame step size and found that the average step size of CENP-A nucleosomes for high salt is double the length of that of low salt (5.5±0.2 nm vs 2.8±0.1 nm, resp., Table 1, Supplemental Figure 5D, Source data 3). Furthermore, we calculated the R step. The R step is the single-frame displacement in the plane of the trajectory and is defined as the square root of the sum of the squares of the displacement in the x and y directions. The maximum R step was only slightly different between the low and high salt conditions (9.6±0.4 nm vs 14.5±0.4 nm, resp., Table 1, Supplemental Figure 6E, Source data 3). When we looked at the total range of a trajectory by calculating the R step range, we observed a much larger variance in the distribution for the high salt compared to the low salt (39.9±2.0 nm^2^ vs 18.3±0.9 nm^2^, resp., Table 1, Supplemental Figure 5F, Source data 3). These results are in line with how salt concentration is known to impact chromatin dynamics (Allahverdi et al., 2015; Brasch et al., 1971; Yager et al., 1989; Yager and Van Holde, 1984).

Next, we plotted the distributions of single step displacements in the x-axis and y-axis and fitted them to a single Gaussian distribution, which is expected for a diffusive process. Whereas the x- and y- single-step distributions were poorly fit by single Gaussian distributions (D0), they were well-fit by a sum of two Gaussian distributions (D1 and D2) (Figure 3D, Supplemental Figure 6). The relative fraction of the two Gaussian distributions differed between low and high salt conditions (Figure 3E, Supplemental Figure 6). These data indicate that there are two distinct populations of nucleosomes in both low and high salt conditions. The D1 Gaussian population represent slower diffusing nucleosomes, potentially reflecting transient pausing, whereas the D2 Gaussian population might represent faster diffusing nucleosomes. The average step distribution (Supplemental Figure 5D) and R step range (Supplemental Figure 5F) display a continuum of data points, instead of a bimodal distribution. This suggests that individual nucleosomes may have the capacity to move back and forth between the D1 and the D2 Gaussian distributions.

Next, we expanded our analyses of the HS-AFM movies of the no APS and 2x APS conditions to MSD, and Gaussian fitting of the single step distributions. As APS functionalization positively charges the mica surface on which chromatin is deposited for HS-AFM imaging, we predicted that CENP-A nucleosomes trajectories would display faster dynamics in the no APS condition vs 2x APS condition. Indeed, the MSD curve of in the absence of APS was higher compared to 2x APS (Figure 3A, Supplementary Figure 4A, B). The average diffusion constant was also higher in the absence of APS (7.5±0.5 nm^2^·s^-1^) than 2x APS (2.5±0.3 nm^2^·s^-1^, Figure 3A, Table 1, Supplemental Figure 5C, Source data 3). A similar pattern was observed for the average step size (6 ±2 nm vs 3.7±0.2 nm, resp.), maximum R step (15.9±0.4 nm vs 11.3±0.5 nm, resp.), and R step variance (43±2 nm^2^ vs 21±1 nm^2^, resp., Table 1, Supplemental Figure 5D-F, Source data 3), with higher values for no APS than 2x APS. The single step displacement distributions were well-fit by a sum of Gaussians (Supplemental Figure 6).

As an additional control, we used a very low concentration of Tween-20 (0.01%), a polysorbate surfactant both helps stabilize proteins and reduces non-specific hydrophobic interactions with the surface. We were interested to learn whether CENP-A chromatin in the presence of Tween- 20 either would display more restricted nucleosomes mobility due to protein stabilization, or less restricted nucleosome mobility due to reduced non-specific interactions. HS-AFM movies of CENP-A chromatin in physiological buffer (0.5x PBS, 2 mM MgCl2) with Tween-20 displayed the most amount of drift (Supplemental Movie 5). After drift correction and filtering, we analyzed the trajectories to obtain single step distributions, MSD curves, and diffusion constants. We found that CENP-A chromatin in the presence of Tween-20 behaved more like no APS and high salt conditions with a steep MSD curve, a very broad distribution of average step sizes, and a large R step range distribution (Figure 3, Supplemental Figure 5, 6, Source data 3). We interpret this to mean that Tween-20 in the context of imaging CENP-A chromatin by HS-AFM primarily reduces non-specific hydrophobic interactions resulting in less restricted nucleosome mobility.

When we analyzed previously published HS-AFM movies of H3 chromatin with or without linker histone H1.5 (Melters and Dalal, 2021) as well as H3 mononucleosomes, we observed a bias in both the angle between successive steps and Gaussian fitting of single step displacements (Supplemental Figure 7, 8, Supplemental movies 6-8, Source data 2, 3). These data provide evidence that bias in HS-AFM trajectories is a possibility and that it can be detected.

Furthermore, there was no difference in fitting either a single or double Gaussian distributions for H3 mononucleosomes (Supplemental Figure 8F). Mononucleosomes are not associated with other nucleosomes and a priori mononucleosomes cannot display whole chromatin fiber motions, allowing mononucleosomes to move freely on the DNA. Therefore, the latter observation implies that the unconstrained motion of mononucleosomes results in a single Gaussian distribution of step displacements.

Altogether, when we analyzed HS-AFM movies, CENP-A chromatin responded to varying salt and APS concentrations in agreement with previous reports (Allahverdi et al., 2015; Brasch et al., 1971; Lyubchenko et al., 2014; Shlyakhtenko et al., 2003; Yager et al., 1989; Yager and Van Holde, 1984). Low salt and 2x APS concentrations reduced CENP-A nucleosome mobility, whereas high salt and no APS concentrations increased CENP-A nucleosome mobility (Supplemental Figure 9). For the remainder of the HS-AFM experiments, we used near physiological relevant salt concentrations (67.5 mM NaCl, 2 mM MgCl2) and standardized APS concentrations (167 nM (Lyubchenko et al., 2014)).

### CENP-C^CD^ represses CENP-A nucleosome mobility *in vitro*

Previously, we showed that a fragment of CENP-C rigidified CENP-A nucleosomes (Melters et al., 2019). Based on these observations, we hypothesized that CENP-C^CD^ would reduce CENP-A nucleosome mobility. To test this hypothesis, we imaged CENP-A chromatin by HS-AFM under near physiological conditions. Subsequently, nucleosome tracks were extracted in either the absence (Figure 4A, Supplemental Figure S1, Supplemental movie 9, Source Data 2, 3) or presence of CENP-C^CD^ (Figure 4B, Supplemental Figure 10, Supplemental movie 10, Source Data 2, 3). From at least 3 experiments per sample, we obtained 498 and 310 trajectories, respectively (Table 1). First, we verified that there were no motion artifacts associated with the tip scanning (Figure 4C, D). Next, we calculated the MSD of CENP-A nucleosomes alone or in the presence of CENP-C^CD^ and found that individual MSD curves of CENP-A nucleosomes were broadly distributed (Figure 4E, Supplemental Figure 10A, B), with an average MSD curve that reached a plateau after ∼25 seconds (Supplemental Figure 10A), implying confined motion (Kapanidis et al., 2018; Zhong and Wang, 2020).

**Figure 4.**
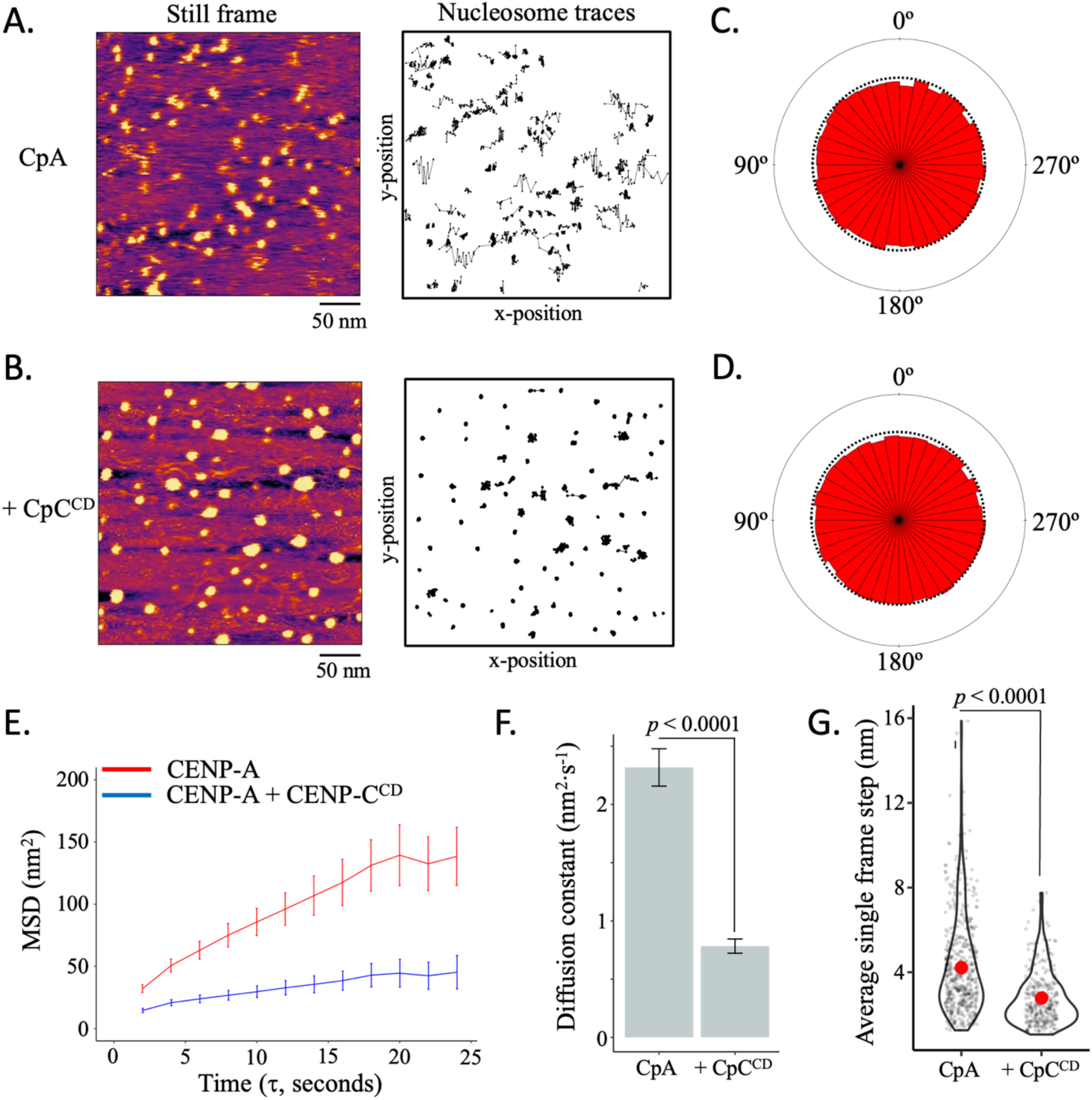
CENP-C^CD^ restrict CENP-A nucleosome mobility *in vitro*. (A) CENP-A nucleosome arrays were tracked in fluid for up to 120 seconds by HS-AFM at 1 frame every 2 seconds. A representative still frame is shown as well as the trajectories over time. (B) CENP-A nucleosome arrays were tracked in the presence of CENP-C^CD^. A representative still frame is shown as well as the trajectories over time. (C) The radial distribution of the angle between successive frames shows a random distribution for CENP-A nucleosomes. (D) The radial distribution of the angle between successive frames shows a random distribution for CENP-A nucleosomes in the presence of CENP-C^CD^. (E) The average mean square displacement is shown with standard error as a function of the time interval. (F) The diffusion constants obtained from the MSD curves. (G) The average single frame step-size for each individual tracked nucleosome (grey points). Red dots represent the overall mean (two-sided t-test; significance was determined at *p*<0.05). The line and bar graphs represent three independent technical replicates.

In contrast, when CENP-C^CD^ was added at 2.2 molar excess to CENP-A nucleosomes (∼one CENP-C^CD^ molecule per CENP-A molecule), qualitatively CENP-A nucleosome mobility was strongly restricted (Figure 4B, Supplemental Figure 10A, B, Supplemental movie 10, Source data 2, 3). Indeed, the MSD curve was much lower compared to CENP-A alone, and the CENP- C^CD^ MSD curve maintained a shallow slope (Figure 4E, Supplemental Figure 10A). The diffusion constant of CENP-A alone was 2.3±0.2 nm^2^·s^-1^, whereas the addition of CENP-C^CD^ reduced the diffusion constant 4-fold to 0.78±0.06 nm^2^·s^-1^ (Figure 4F, Table 1, Supplemental Figure 10C, Source Data 2, 3). This difference is reflected in the smaller single frame step size when CENP-C^CD^ is added to CENP-A chromatin compared to CENP-A chromatin alone (Figure 4G, Supplemental Figure 10D, Source data 3). The maximum R step and R step range were also larger for CENP-A nucleosomes without CENP-C^CD^ (Supplemental Figure 10E, F, Source data 3). Next, we fitted the single step displacement distributions with Gaussian distributions and found that a sum of two Gaussian distributions provided better fits to the single step displacement distributions (Supplement Figure 11). Overall, by HS-AFM we observed that CENP-C^CD^ restricts CENP-A nucleosome mobility. This, we hypothesized could be the mechanism by which overexpression of CENP-C in living cells results in CENP-A chromatin clustering, as we recently observed (Melters et al., 2019).

### Excess CENP-C suppresses centromeric chromatin accessibility and transcription *in vivo*

Next, we asked what the functional consequences are of CENP-C on CENP-A chromatin, beyond the formation of kinetochores (Cheeseman et al., 2006; Kai et al., 2021; Mendiburo et al., 2011; Régnier et al., 2005). Previously we showed that overexpressing CENP-C in HeLa cells resulted in increased clustering of centromeric chromatin and loss of centromeric RNAP2 (Melters et al., 2019). These results, combined with our “loss of motion” observations above, suggest that centromeric noncoding α-satellite transcription might be impaired in the background of CENP-C overexpression. To examine this facet of CENP-C:CENP-A homeostasis, we overexpressed CENP-C in HeLa cells to assess if centromeric transcription is altered.

First, we measured the effects of CENP-C overexpression on the level of RNAP2 on CENP-A chromatin. We first pulled-down CENP-A chromatin associated with CENP-C by CENP-C native ChIP (nChIP), and the unbound CENP-A chromatin was pulled-down by sequential ACA nChIP. We found that RNAP2 levels were reduced upon CENP-C overexpression (Figure 5A, Supplemental Figure 11, Source Data 4). In addition, we observed that, upon CENP-C overexpression, total CENP-A levels were also reduced (Figure 5B, Supplemental Figure 12, Source Data 4). Previously (Melters et al., 2019), we showed that the addition of CENP-C^CD^ or overexpression of CENP-C results in compaction of CENP-A chromatin. We therefore wondered if CENP-C overexpression impacted centromeric transcription. By quantitative PCR, we observed a ∼60% reduction in α-satellite transcripts in cells overexpressing CENP-C (Figure 5C, Source data 5).

**Figure 5.**
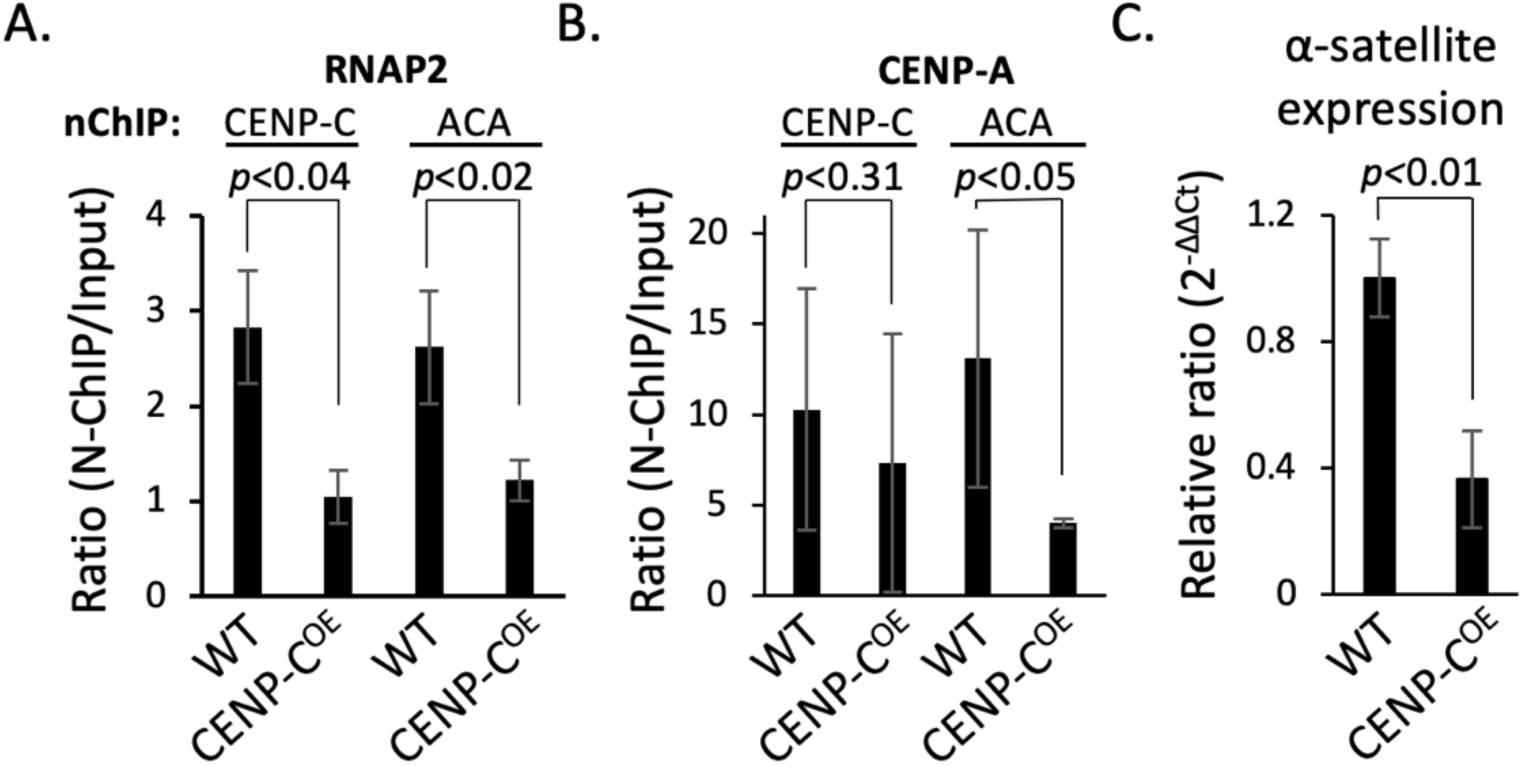
CENP-C overexpression suppressed a-satellite expression and centromeric RNAP2 occupancy. (A) Quantification of RNAP2 levels pulled down with either CENP-C or sequential ACA nChIP. (B) Quantification of CENP-A levels that pulled down with either CENP-C or sequential ACA nChIP. (C) Quantification of consensus α-satellite transcription in mock-transfected (WT) and CENP-C overexpression (CENP-C^OE^) (two-sided t-test; significance was determined at *p*<0.05). The bar graphs represent three independent technical replicates, and the error bars represent standard deviations.

These results indicate that CENP-A levels are reduced and that centromeric transcription is indeed impaired upon CENP-C overexpression. Previous reports showed that new CENP-A loading is transcriptionally regulated (Jansen et al., 2007; Quénet and Dalal, 2014a). Therefore, we hypothesized that CENP-C overexpression leads to defective *de novo* CENP-A loading.

### CENP-C overexpression limits *de novo* CENP-A loading

To test this hypothesis, we turned to the well-established SNAP-tagged CENP-A system combined with quench pulse-chase immunofluorescence (Bodor et al., 2012). Using this system in cells synchronized to mid-G1, one can distinguish between older CENP-A (TMR-block) and newly incorporated CENP-A (TMR-Star) (Figures 6A, B). Strikingly, in an CENP-C overexpression background, we observed a 2.3-fold reduction of *de novo* incorporation of CENP-A (Figures 6C, D, Source Data 6).

**Figure 6.**
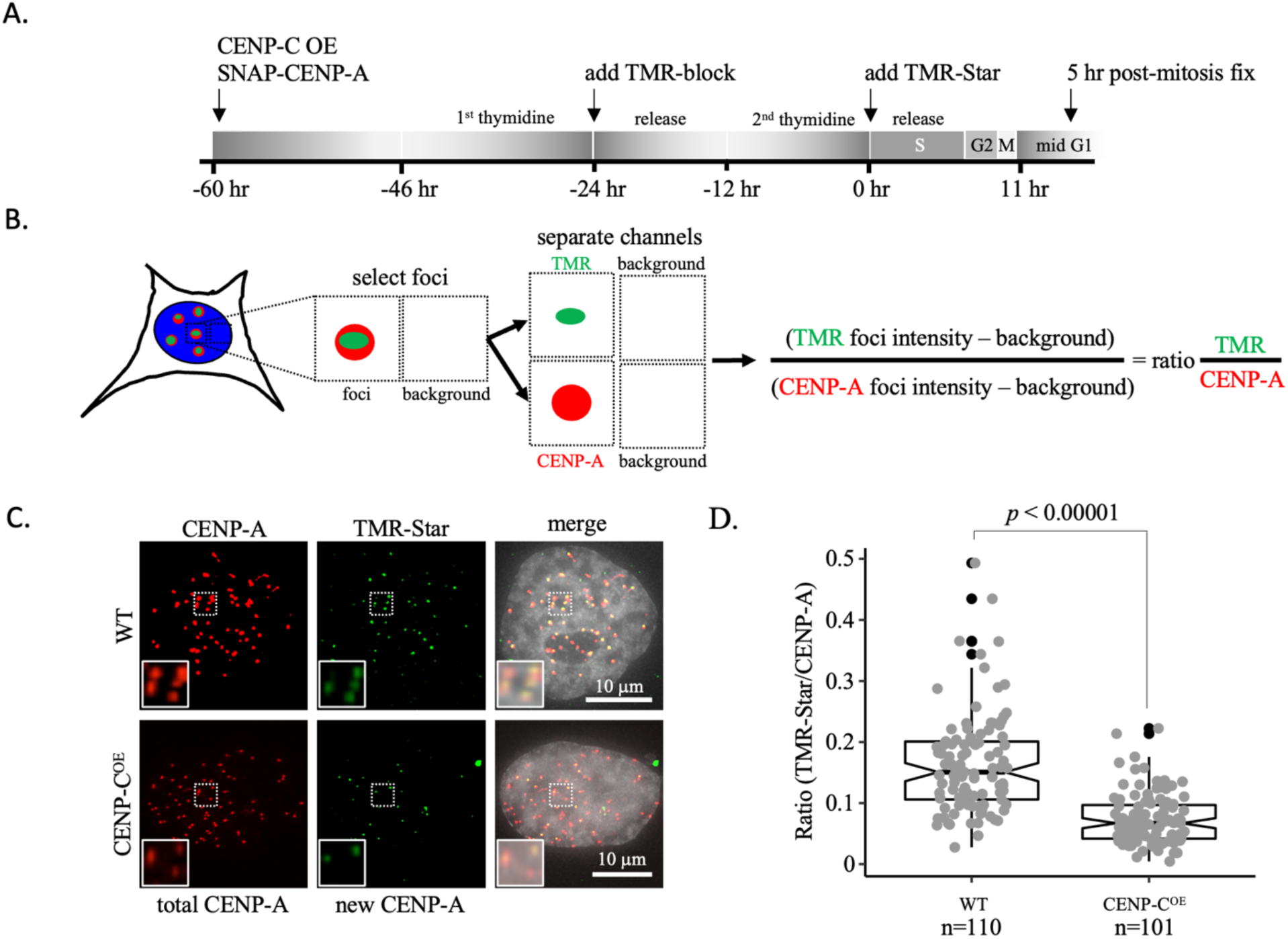
New CENP-A loading impaired upon CENP-C overexpression. (A) Schematic of experimental design. (B) Colocalized immunofluorescent signals for CENP-A and TMR-Star are collected and the intensity of both foci is measured as well as background directly neighboring the foci to determine the ratio of the TMR-star signal over total CENP-A signal. (C) *De novo* CENP-A incorporation was assessed by quench pulse-chase immunofluorescence. After old CENP-A was quenched with TMR-block, newly loaded CENP-A was stained with TMR-Star and foci intensity was measured over total CENP-A foci intensity. Inset is a 2x magnification of the dotted box in each respective image. (D) Quantification of *de novo* CENP-A loading by measuring the ratio of TMR-Star signal over total CENP-A signal (one-way ANOVA; significance was determined at *p*<0.05). The box plots represent three independent technical replicates.

Thus, the functional consequences of CENP-C overexpression are suppression of α-satellite transcription (Figure 5C) and subsequent impairment of new CENP-A loading (Figures 6C, D).

## Discussion

Here, we demonstrate that HS-AFM is a reliable technique for studying single nucleosome dynamics by directly visualizing their motion in real time. Importantly, we show that the AFM tip scanning motion does not generate observable artifacts in the nucleosome dynamics (Figure 2, Supplemental Figure 3, 4). Various critical technical conditions were tested to assess their impact on CENP-A chromatin dynamics. Salt is a well-known to either stabilize or destabilize chromatin (Allahverdi et al., 2015; Brasch et al., 1971; Lyubchenko et al., 2014; Shlyakhtenko et al., 2003; Yager et al., 1989; Yager and Van Holde, 1984), as salt dialysis is a common method for *in vitro* reconstitution of nucleosomes (Cruz-Becerra and Kadonaga, 2021; Peterson, 2008; Walkiewicz et al., 2014). Indeed, low salt restricted CENP-A nucleosome motion, whereas high salt did the opposite (Figure 3, Supplemental Figure 5, 6). In AFM studies, APS is used to functionalize the mica surface with positive charges facilitating the interaction between DNA or chromatin to the surface. Not functionalizing the mica surface resulted in more mobile CENP-A nucleosomes, whereas 2x APS functionalized mica resulted in increased levels of immobile CENP-A nucleosomes (Figure 3, Supplemental Figure 5, 6). In addition, H3 mononucleosomes were the only nucleosomes for which the single-frame step size distribution was well-fit by a single Gaussian distribution (Supplemental Figure 8F). Altogether, these data provide evidence that CENP-A chromatin responds to both buffer conditions and surface functionalization in a predictable manner.

CENP-C modulates both CENP-A nucleosome accessibility (Ali-Ahmad et al., 2019; Ariyoshi et al., 2021; Falk et al., 2016, 2015; Guo et al., 2017) and its elasticity (Melters et al., 2019). Here, we show that CENP-C^CD^ regulates CENP-A nucleosomes mobility as well (Figure 4, Supplemental Figure 10, 11). The data presented here correlates with CENP-C^CD^ rigidification of CENP-A nucleosomes (Melters et al., 2019). *In vivo*, CENP-C overexpression results in reduced levels of centromeric RNAP2 (Melters et al., 2019), impaired centromeric transcription (Figure 5C), and subsequent decreased loading of new CENP-A at the centromere (Figure 6). These findings suggest that physical properties of nucleosomes dictate their mobility along DNA, which in turn regulates chromatin accessibility and transcriptional potential.

In the nucleus, nucleosomes move in different dimensions, either in a single dimension along the DNA strand, or three dimensions where the DNA strand moves and the nucleosomes follow as passengers (Babokhov et al., 2020; Ide et al., 2022; Melters and Dalal, 2021). In addition, various events, ranging from transcription to DNA repair and replication, involve chromatin remodeling (Nodelman and Bowman, 2021; Zaret, 2020). Several *in vivo* studies have utilized tagged H2B combined with high-resolution live cell imaging to probe nucleosome dynamics.

Nucleosomes within heterochromatin were less dynamic compared to euchromatin regions (Nozaki et al., 2017; Ricci et al., 2015) and local levels of H1 correlated with reduced nucleosome dynamics (Gómez-García et al., 2021). Studies using optical tweezers have probed the one dimensional diffusive behavior of nucleosomes (Chen et al., 2019; Rudnizky et al., 2019). A recent study using optical tweezers showed that the diffusion constant of H3 mononucleosomes sliding along a DNA strand is ∼1.3 bp^2^·s^-1^ (Rudnizky et al., 2019) (∼0.15 nm^2^·s^-1^). Binding of transcription factors to nucleosomal DNA biases nucleosome motion, potentially through manipulation of the folding and unfolding of nucleosomal DNA around the nucleosome core particle (Donovan et al., 2023; Rudnizky et al., 2019). Here, we show that CENP-A nucleosomes have a diffusion constant of 2.3±0.2 nm^2^·s^-1^ and was well-fit by a double Gaussian distribution (Figure 4, Supplemental Figure 11). When CENP-C^CD^ was added at a 2.2 molar ratio to CENP-A nucleosomes, we observed a diffusion constant that is almost 3-times lower than CENP-A nucleosomes alone (0.78 ± 0.06 nm^2^·s^-1^). This difference in diffusion constants for nucleosomes between the Rudnizky study (∼0.15 nm^2^·s^-1^, (Rudnizky et al., 2019)) and our results (0.78-2.32 nm^2^·s^-1^) could be due to technical differences of their experimental set-up compared to our experimental approach (optical tweezers vs HS-AFM). We used nucleosome arrays which permit both one-dimensional (nucleosome sliding) and three-dimensional motions (whole chromatin fiber movement) of nucleosomes compared to mononucleosomes that were fixed to polystyrene beads, which only permit one-dimensional motion (Rudnizky et al., 2019). In contrast, single molecule tracking in cultured cells captures nucleosome dynamics within the nucleus (Iida et al., n.d.; Kimura and Cook, 2001; Morisaki et al., 2014), but requires fluorophore-tags. Fluorophore-tags are photosensitive (Jradi and Lavis, 2019) and have the potential of functionally altering the function of the protein it is bound to (Maheshwari et al., 2015; Ravi et al., 2010). Calculated diffusion constants range for H2B-eGFP range from 0.0019 µm^2^·s^-1^ to 7.3 µm^2^·s^-1^ (Bhattacharya et al., 2006; Mazza et al., 2012) and 0.03 µm^2^·s^-1^ to 1.39 µm^2^·s^-1^ for HALO-H2B (Lovely et al., 2020; Ranjan et al., 2020). These *in vivo* H2B diffusion constants are mostly several order of magnitudes larger than what we observed (0.78-2.32 nm^2^·s^-1^). This could be due to the many activities within the nucleus of a living cell compared to the steady-state nature of an *in vitro* experiment. As such, the potential of single molecule analysis by HS-AFM to contribute to elegant studies in the field is exciting, as it can link single molecule force spectroscopy analysis to *in vivo* single molecule tracking experiments.

Looking at another histone variant nucleosome, the diffusion constant of H2A.Z nucleosomes is larger than H3 nucleosomes (Rudnizky et al., 2019), which correlates with the transcriptional buffering function of H2A.Z (Chen et al., 2019; Giaimo et al., 2019). A logical prediction from our results would be that transcription through CENP-A chromatin would be more efficient compared to H3 chromatin. Indeed, a recent single-molecule study showed that CENP-A creates an open chromatin structure (Nagpal and Fierz, 2022). We found that proteins that exert their activity by binding to the outer surface of nucleosomes, rather than changing the internal constituents of nucleosomes, have the opposite effect on transcription. CENP-C^CD^ rigidifies CENP-A nucleosomes (Melters et al., 2019), and CENP-C^CD^-CENP-A nucleosomes have a significantly reduced diffusion constant compared to CENP-A nucleosomes alone (Table 1). *In vivo*, overexpression of CENP-C resulted in decreased centromeric transcription (Figure 6).

Thus, this study significantly extends the existing paradigm that suggests nucleosome dynamics is a highly tunable platform, with a surprising twist - tuners can dampen or exaggerate nucleosome motion, which correlates with higher order folding and accessibility.

Taking these and prior findings in context, our data suggest a model where chromatin effector partners modify the material properties of histone variant nucleosomes at the local level, supporting the formation of functional nucleosome clutches (Portillo-Ledesma et al., 2021; Ricci et al., 2015) (Figure 7). Here, we specifically probed CENP-A chromatin and based on our results we propose the following model. CENP-A nucleosomes recruit CENP-C to form a kinetochore-promoting CENP-A chromatin clutch. CENP-C functions as the blueprint for the recruitment of additional kinetochore proteins (Klare et al., 2015). When CENP-C binds, CENP-A nucleosomes become rigidified (Melters et al., 2019) and restrict CENP-A nucleosomes mobility (Figure 4). This unique clutch facilitates the recruitment of other inner kinetochore components by locally immobilizing CENP-A chromatin. Yet, to load new CENP-A molecules, transcription must happen. The repressive chromatin state that CENP-C induces contradicts this functional necessity. One speculative manner in which cells can work around the juxtaposition of the dual functions of CENP-A chromatin is by maintaining a pool of free centromeric CENP-A nucleosomes (Figure 7). Indeed, ChIP-seq and FISH data have established that centromeric CENP-C levels are lower compared to CENP-A levels (Henikoff et al., 2015; Kyriacou and Heun, 2018). We propose that unbound elastic and mobile CENP-A chromatin clutch create an intrinsically accessible chromatin state, allowing for the recruitment of transcriptional machinery that maintains centromeric CENP-A levels to facilitate both opposing functions (Figure 7).

**Figure 7.**
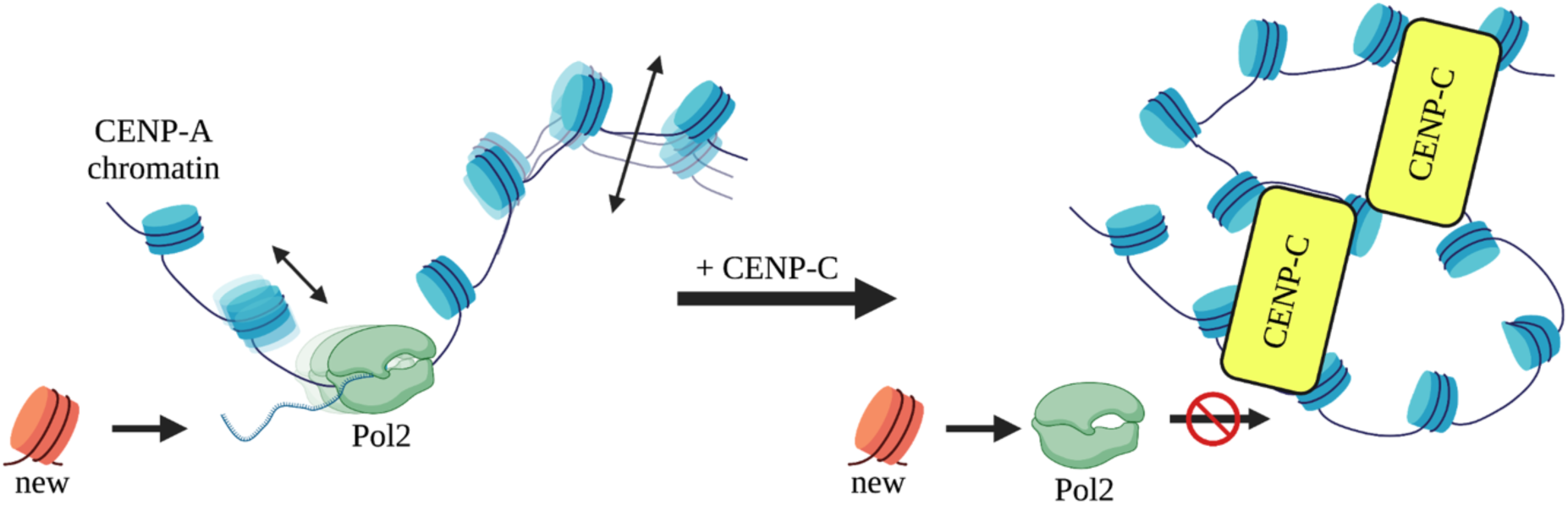
Clutch model for CENP-C restricted motion of CENP-A chromatin. Under wildtype conditions, we propose that CENP-A chromatin not bound by CENP-C (yellow box) form a chromatin clutch is readily accessible to the transcriptional machinery, because of the intrinsic material properties of CENP-A nucleosomes. In contrast, when CENP-C or CENP-C complexes bind CENP-A nucleosomes, a unique clutch of CENP-A chromatin is formed restricting sliding of CENP-A nucleosomes. This coincides with CENP-C^CD^ altering the material properties of and quenches of CENP-A’s mobility. Less mobile CENP-A nucleosomes restrict progression of the transcriptional machinery and subsequent loading of new CENP-A molecules.

Succinctly, the epigenetic fate of a locus may be tightly, possibly causally, linked to the mechanical state of the chromatin fiber; concomitantly, the reinstatement of an epigenetic signature by *de novo* loading of a particular variant, reinforces the mechanical state of the fiber.

In summary, we report the MSD and diffusion constant for CENP-A nucleosomes and is well-fit by a double Gaussian distribution. CENP-C modulates centromeric chromatin accessibility by restricting the motions of CENP-A nucleosomes. *In vivo*, such nucleosome mobility changes correlate with diminished transcriptional competence, resulting in suppression of *de novo* CENP-A loading. Nucleosome dynamics play an important role in genome compaction and regulating DNA access by DNA binding factors. These dynamics are driven by only a few interactions between the interfaces of DNA and nucleosomes (Fierz and Poirier, 2019; Polach and Widom, 1995; Widom, 1998). An exciting line of investigation is to examine how at the nanoscale level the interaction between CENP-A nucleosomes and CENP-C protein co-evolved to repress

CENP-A’s mobile and elastic nature. It will also be crucial to examine how CENP-A:CENP-C homeostasis, along with those of other key inner kinetochore proteins such as CENP-N, T, S, X, and W regulate centromeric transcription and thereby *de novo* assembly of CENP-A to maintain the epigenetic identity of centromeres in other species. At a more global scale, it will be exciting to ask how different H1 variants modulate the chromatin fiber at the local level to promote or limit transcriptional competency, and how H1 variants dictate the mechanical motions of individual nucleosomes within the local 10nm chromatin fiber.

## Material and Methods

### Key Resources Table

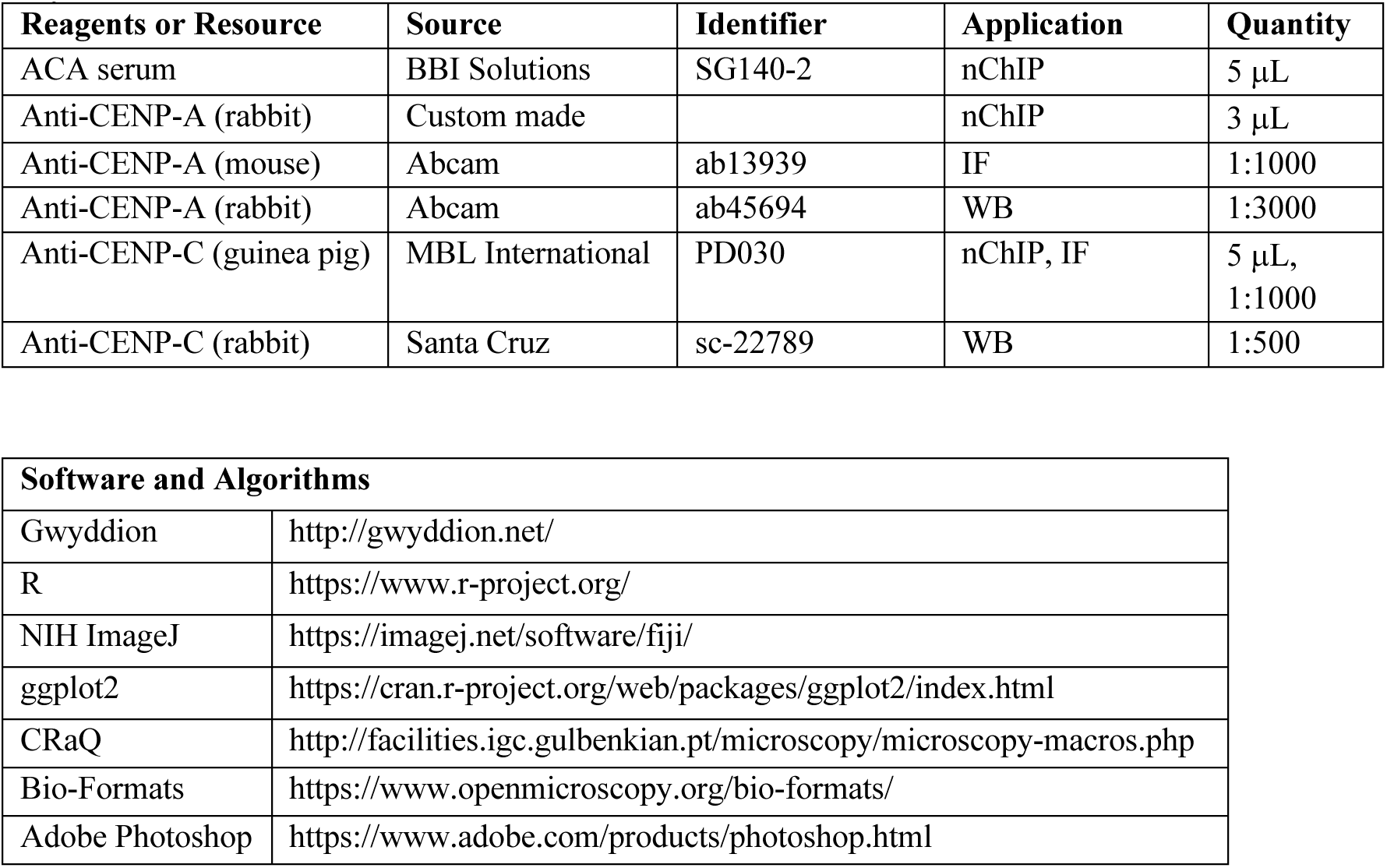

| Reagents or Resource | Source | Identifier | Application | Quantity |
| --- | --- | --- | --- | --- |
| ACA serum | BBI Solutions | SG140-2 | nChIP | 5 μL |
| Anti-CENP-A (rabbit) | Custom made | | nChIP | 3 μL |
| Anti-CENP-A (mouse) | Abcam | ab13939 | IF | 1:1000 |
| Anti-CENP-A (rabbit) | Abcam | ab45694 | WB | 1:3000 |
| Anti-CENP-C (guinea pig) | MBL International | PD030 | nChIP, IF | 5 μL,<br>1:1000 |
| Anti-CENP-C (rabbit) | Santa Cruz | sc-22789 | WB | 1:500 |

| Software and Algorithms |  |
| --- | --- |
| Gwyddion | http://gwyddion.net/ |
| R | https://www.r-project.org/ |
| NIH ImageJ | https://imagej.net/software/fiji/ |
| ggplot2 | https://cran.r-project.org/web/packages/ggplot2/index.html |
| CRaQ | http://facilities.igc.gulbenkian.pt/microscopy/microscopy-macros.php |
| Bio-Formats | https://www.openmicroscopy.org/bio-formats/ |
| Adobe Photoshop | https://www.adobe.com/products/photoshop.html |

### In vitro reconstitution

*In vitro* reconstitution of CENP-A (CENP-A/H4 cat#16-010 and H2A/H2B cat#15-0311, EpiCypher, Research Triangle Park, NC) and H3 (H3/H4 cat#16-0008 and H2A/H2B cat#15-0311, EpiCypher Research Triangle Park, NC) nucleosomes were performed as previously described (Dimitriadis et al., 2010; Walkiewicz et al., 2014). For quality purposes, an aliquot of each sample was imaged by AFM in non-contact tapping mode, before moving on to high-speed AFM.

### High-speed AFM

*In vitro* reconstituted CENP-A and H3 chromatin with the addition of CENP-C^CD^ (Melters et al., 2019) or H1.5 (Melters and Dalal, 2021), resp. was imaged with the Cypher VRS (Oxford Instruments, Asylum Research, Santa Barbara, CA) using ultra-small silicon cantilevers (BL- AC10DS with nominal resonances of ∼1500 kHz, stiffness of ∼0.1 N/m) in non-contact-mode.

The mica on top of the scanning pilar was peeled and functionalized with 30 µM APS, before 10 µL of sample was added. The sample was incubated for 15-20 minutes before initializing scanning to obtain a density of ∼400 nucleosome/µm^2^. The sample was imaged at a speed of 268.82 Hz (frame rate = 0.5Hz or one frame per two second) and a resolution of 512x256 points and lines for an area of 400x400nm (CENP-A and CENP-A + CENP-C^CD^ sample with a nucleosome density of ∼300 nucleosomes/µm^2^, respectively), 300x300 nm (H3 + H1.5 sample with a nucleosome density of ∼200 nucleosomes/µm^2^), and 250x250 nm (H3 sample with a nucleosome density of ∼200 nucleosomes/µm^2^). Movies were saved in mp4 format, converted to TIFF sequences using Photoshop (Adobe), prepared for single molecule tracking in ImageJ (Fuji), and tracked with MATLAB’s single particle tracking package. Obtained nucleosome tracks that were shorter than 10 frames and had single steps exceeding 24 nm were excluded from the analysis. Using ggplot2 package in R (version 4.2.1), drift was visualized and calculated for each individual movie and any observed drift was corrected and verified. The remaining nucleosome tracks were subsequently analyzed to obtain the mean square displacement curves, diffusion constant, angle between successive frames, single frame step sizes, the maximum R step, and R step range. The R-step is the single-frame displacement in the plane. It is defined as the square root of the sum of the squares of the displacement in the x and y direction [R step = 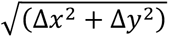]. The results were visualized using the ggplot2 package in R (version 4.2.1). The diffusion constant for each nucleosome sample was estimated from the initial two points of each MSD (Vestergaard et al., 2014). The mean and standard error of the mean are reported as the diffusion constant and uncertainty for each nucleosome sample. The mean diffusion constant is identical to the diffusion constant derived from the first two points of the average MSD curves for each nucleosome sample.

### Native Chromatin-Immunoprecipitation and Western blotting

HeLa cells were grown in DMEM (Invitrogen/ThermoFisher Cat #11965) supplemented with 10% FBS and 1X penicillin and streptomycin cocktail. nChIP experiments were performed without fixation. After cells were grown to ∼80% confluency, they were harvested as described (Bui et al., 2017, 2012). For best results, the pellet obtained for chromatin were spun-down during the nuclei extraction protocol (Walkiewicz et al., 2014) and was broken up with a single gentle tap. Nuclei were digested for 6 minutes with 0.25 U MNase/mL (Sigma-Aldrich cat #N3755-500UN) and supplemented with 1.5 mM CaCl2. Following quenching (10 mM EGTA), nuclei pellets were spun down, and chromatin was extracted gently, overnight in an end-over-end rotator, in low salt solution (0.5x PBS; 0.1 mM EGTA; protease inhibitor cocktail (Roche cat #05056489001). nChIP chromatin bound to Protein G Sepharose beads (GE Healthcare cat #17- 0618-02) were gently washed twice with ice cold 0.5x PBS and spun down for 1 minute at 4°C at 800 rpm. Following the first nChIP, the unbound fraction was used for the sequential nChIP. Western analyses were done using LiCor’s Odyssey CLx scanner and Image Studio v2.0.

### Quantitative PCR

α-satellite expression levels in HeLa cells that were either mock transfected or transfected GFP-CENP-C were detected as previously described (McNulty et al., 2017; Quénet and Dalal, 2014b). RNA was extracted, quantified by UV-spectroscopy, and equal quantities were retro-transcribed using Superscript III First-Strand Synthesis kit as described above. Complementary DNA (cDNA) samples were prepared using the iQ SYBR Green supermix (#170–8880; Biorad) following manufacturer’s protocol. Control reactions without cDNA were performed to rule out non-specific amplification. The qPCR was run on Step one plus Real time PCR system (Applied Biosystem, Grand Island, NY). Primer sequences are:

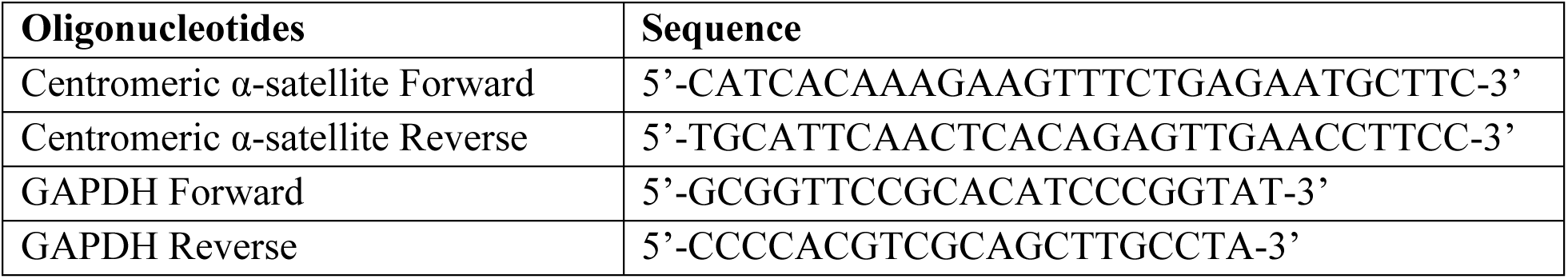

The comparative cycle threshold (CT) method was used to analyze the expression level of α- satellite transcripts. CT values were normalized against the average CT value of the housekeeping gene GAPDH. Relative fold differences (2^−ΔΔCT^) are indicated in Figure 5C.

### Quench pulse-chase immunofluorescence

To quantify *de novo* assembled CENP-A particles, we transfected HeLa cells with SNAP-tagged CENP-A (generous gift from Dan Foltz) in combination with either empty vector or GFP-CENP-C using the Amaxa Nucleofector kit R (Lonza Bioscience, Walkersville, MD) per instructions.

The quench pulse-chase experiment was performed according to Bodor et al [42]. In short, following transfection, cells were synchronized with double thymidine block. At the first release TMR-block (S9106S, New England Biolabs, Ipswich, MA) was added per manufactures instruction and incubated for 30 min at 37°C, followed by three washes with cell culture media. At the second release TMR-Star (S9105S, New England Biolabs, Ipswich, MA) was added per manufactures instructions and incubated for 15 min at 37°C, followed by three washes with cell culture media. Fourteen hours after adding TMR-Star, cells were fixed with 1% paraformaldehyde in PEM (80 mM K-PIPES pH 6.8, 5 mM EGTA pH 7.0, 2 mM MgCl_2_) for 10 min at RT. Next, cells were washed three times with ice cold PEM. To extract soluble proteins, cells were incubated with 0.5% Triton-X in CSK (10 mM K-PIPES pH 6.8, 100 mM NaCl, 300 mM sucrose, 3 mM MgCl2, 1 mM EGTA) for 5 min at 4°C. The cells were rinsed with PEM and fixed for a second time with 4% PFA in PEM for 20 min at 4°C. Next, the cells were washed three times with PEM. Cells were permeabilized with 0.5% Triton-X in PEM for 5 min at RT and subsequently washed three times with PEM. Next, the cells were incubated in blocking solution (1X PBS, 3% BSA, 5% normal goat serum) for 1 hr at 4°C. CENP-A antibody (ab13979 1:1000) was added for 1 hr at 4°C, followed by three washes with 1X PBS-T. Anti-mouse secondary (Alexa-488 1:1000) was added for 1hr at 4°C, followed by three 1X PBS-T and two 1X PBS washes. Following air-drying, cells were mounted with Vectashield with DAPI (H- 1200, Vector Laboratories, Burlingame, CA) and the coverslips were sealed with nail polish.

Images were collected using a DeltaVision RT system fitted with a CoolSnap charged-coupled device camera and mounted on an Olympus IX70. Deconvolved IF images were processed using ImageJ. From up to 22 nuclei, colocalizing CENP-A and TMR-Star foci signal were collected, as well as directly neighboring regions. Background signal intensity was subtracted from corresponding CENP-A and TMR-Star signal intensity before the ratio CENP-A/TMR-Star was determined. Graphs were prepared using the ggplot2 package for R.

### Quantification and statistical analyses

Significant differences for Western blot quantification and nucleosome track measurements from HS-AFM analyses were performed using either paired or two-sided t-test or one-way ANOVA as described in the figure legends. Significance was determined at p<0.05.

## Acknowledgements

We thank Drs. Tom Misteli and Sam John, and members of the CSEM laboratory for critical comments and suggestions. We thank Drs. Ankita Saha and Craig Mizzen (deceased, University of Illinois, Urbana-Champaign) for the gift of recombinant H1.5 protein. We thank Dr. Will Heinz for kindly letting us use his custom set-up for HS-AFM. We thank the reviewers for very useful feedback. This work was supported by the intramural research fund of the Center for Cancer Research at the National Cancer Institute/NIH (D.P.M., T.R., and Y.D.) and National Heart, Lung, and Blood Institute (K.C.N.).

## Supplemental Figures

**Supplemental Figure 1.**
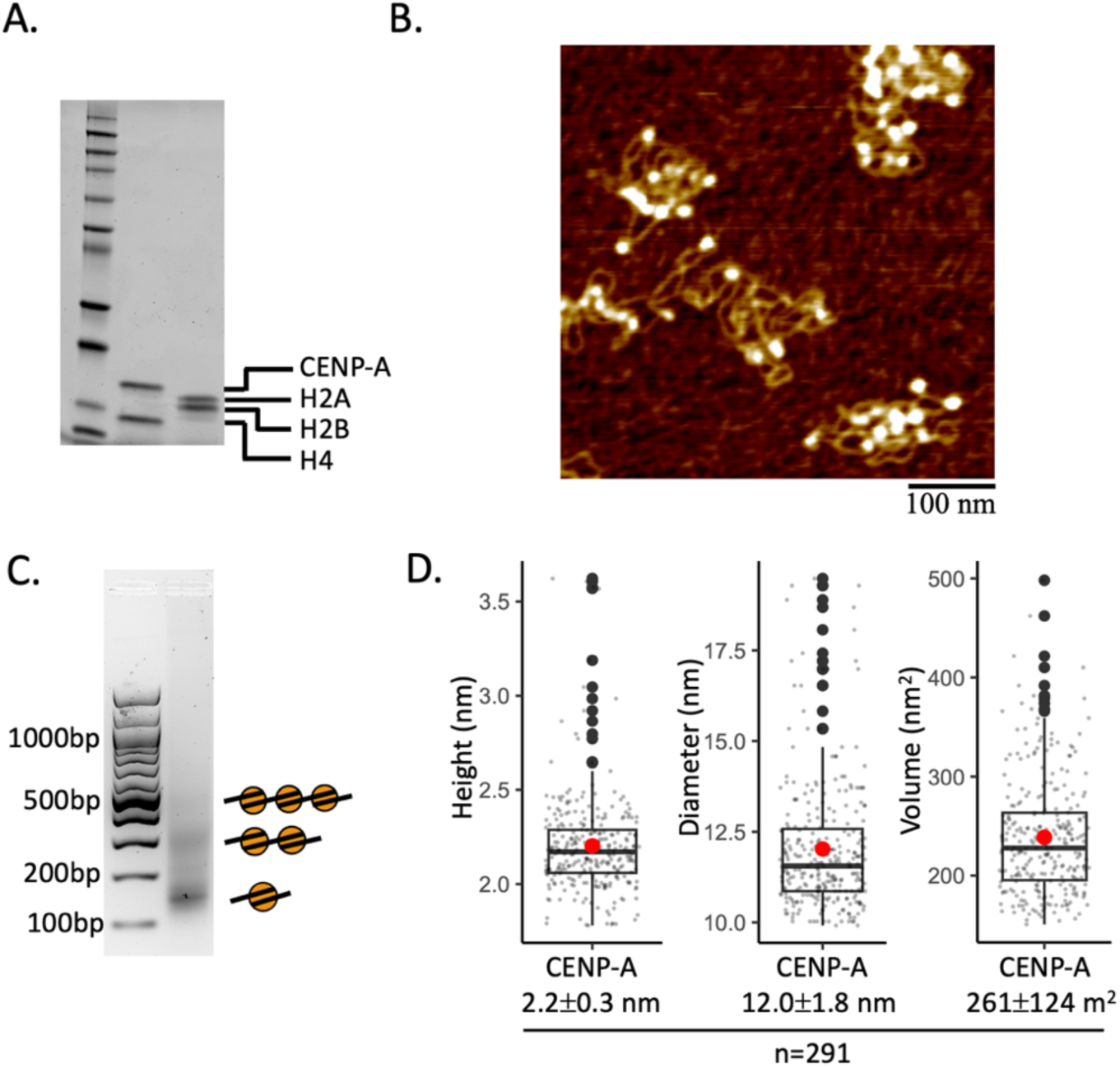
(A) Purified histone protein was quantified from SDS-PAGE blots for calculating the correct *in vitro* reconstitution ratios. (B) Representative in air AFM image of *in vitro* reconstituted CENP-A chromatin. (C) Agarose gel of MNase digested *in vitro* reconstituted CENP-A chromatin showing mono-, di-, and trinucleosome arrays confirming chromatin formation. (D) Quantification of nucleosome measurements of *in vitro* reconstituted CENP-A nucleosome arrays.

**Supplemental Figure 2.**
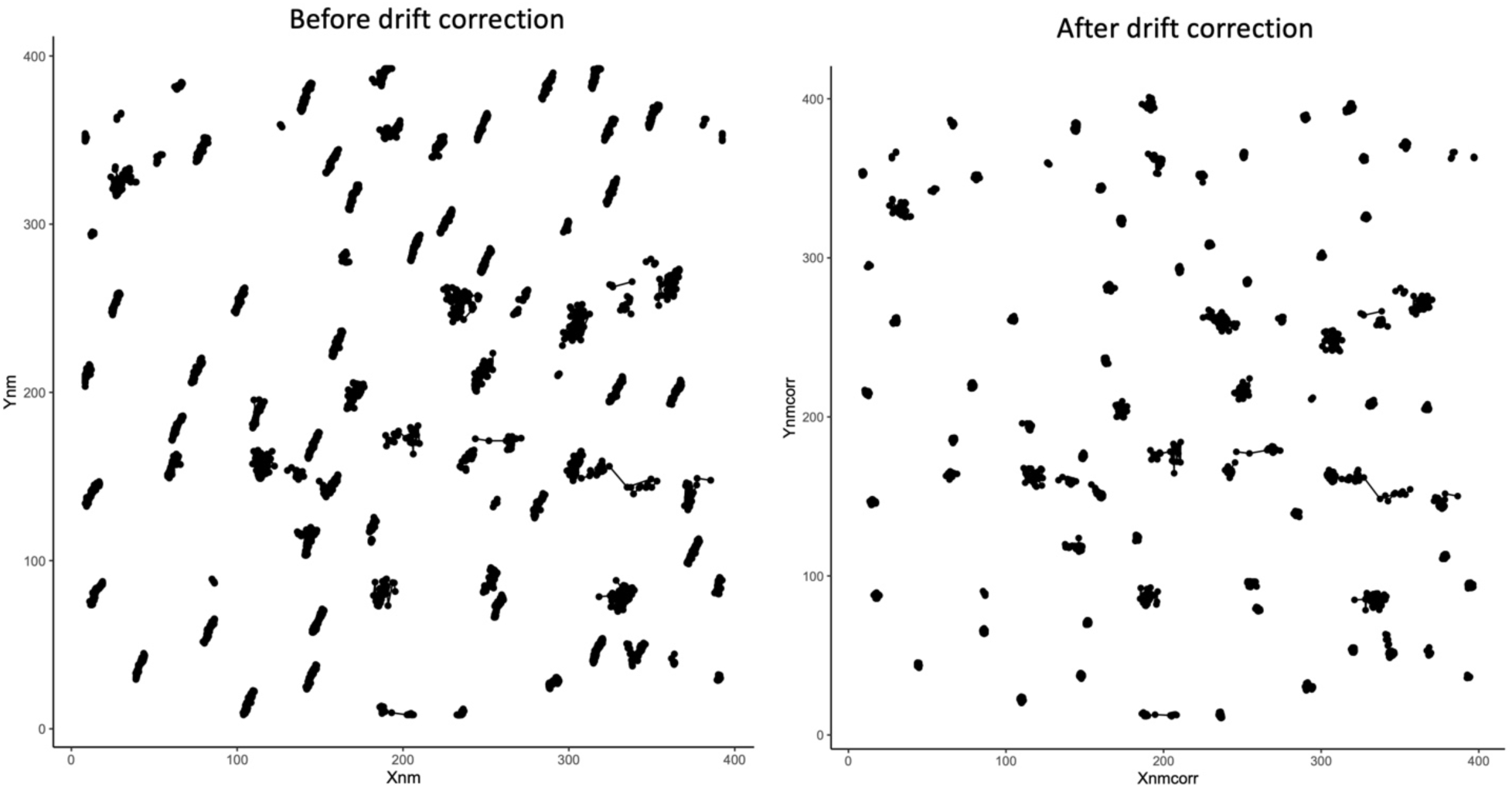
A representative example drift correction. Trajectories that do not appear to move much but do show a streak were used to calculate and correct for drift. After drift corrections, the results were verified.

**Supplemental Figure 3.**
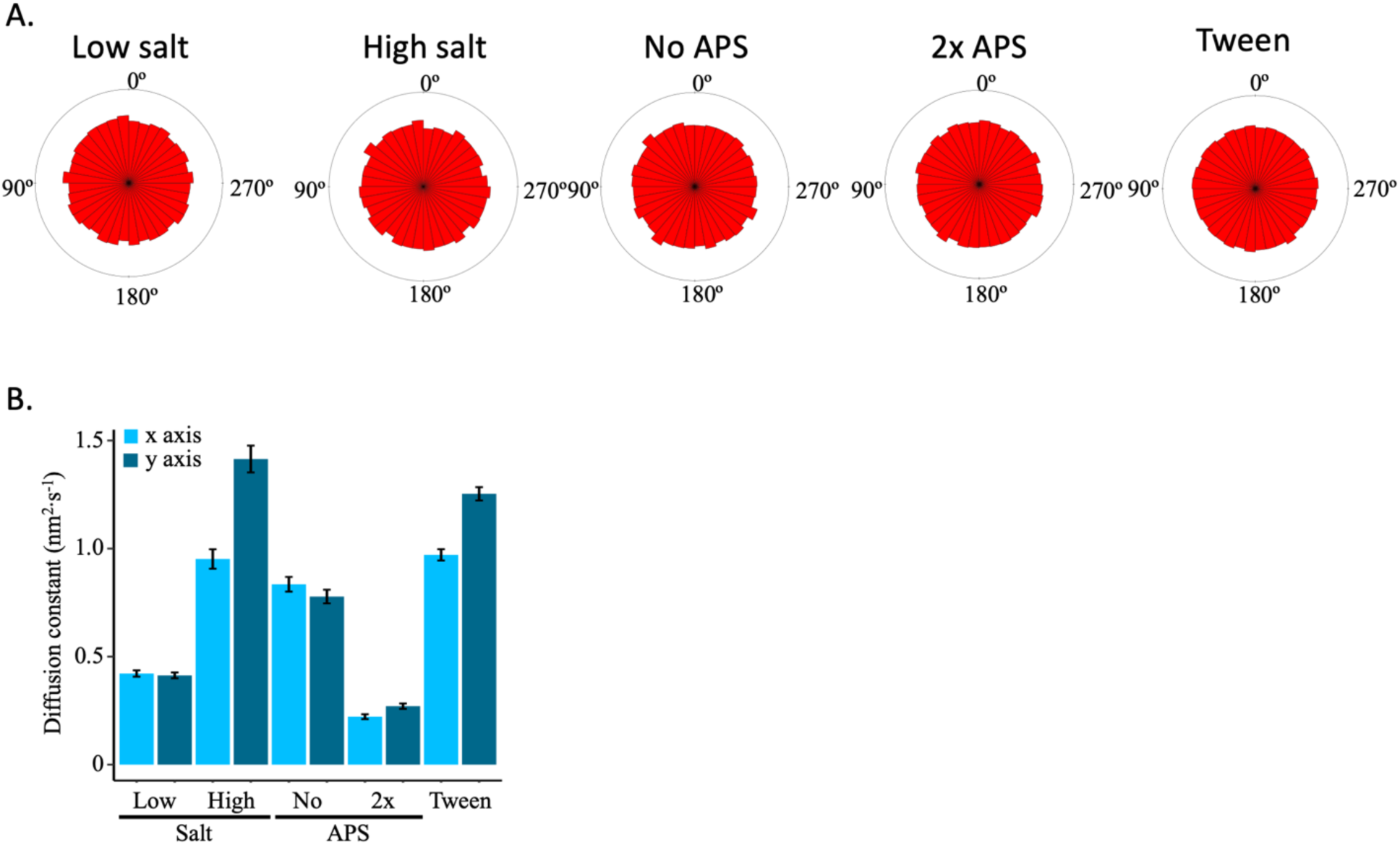
(A) The distribution of the angle between successive nucleosome positions for each control condition. (B) Diffusion constants estimated from the x- and y-axis displacement distributions for each individual control condition. The data is obtained from two independent technical replicates per condition. The error bars represent standard error of the mean.

**Supplemental Figure 4.**
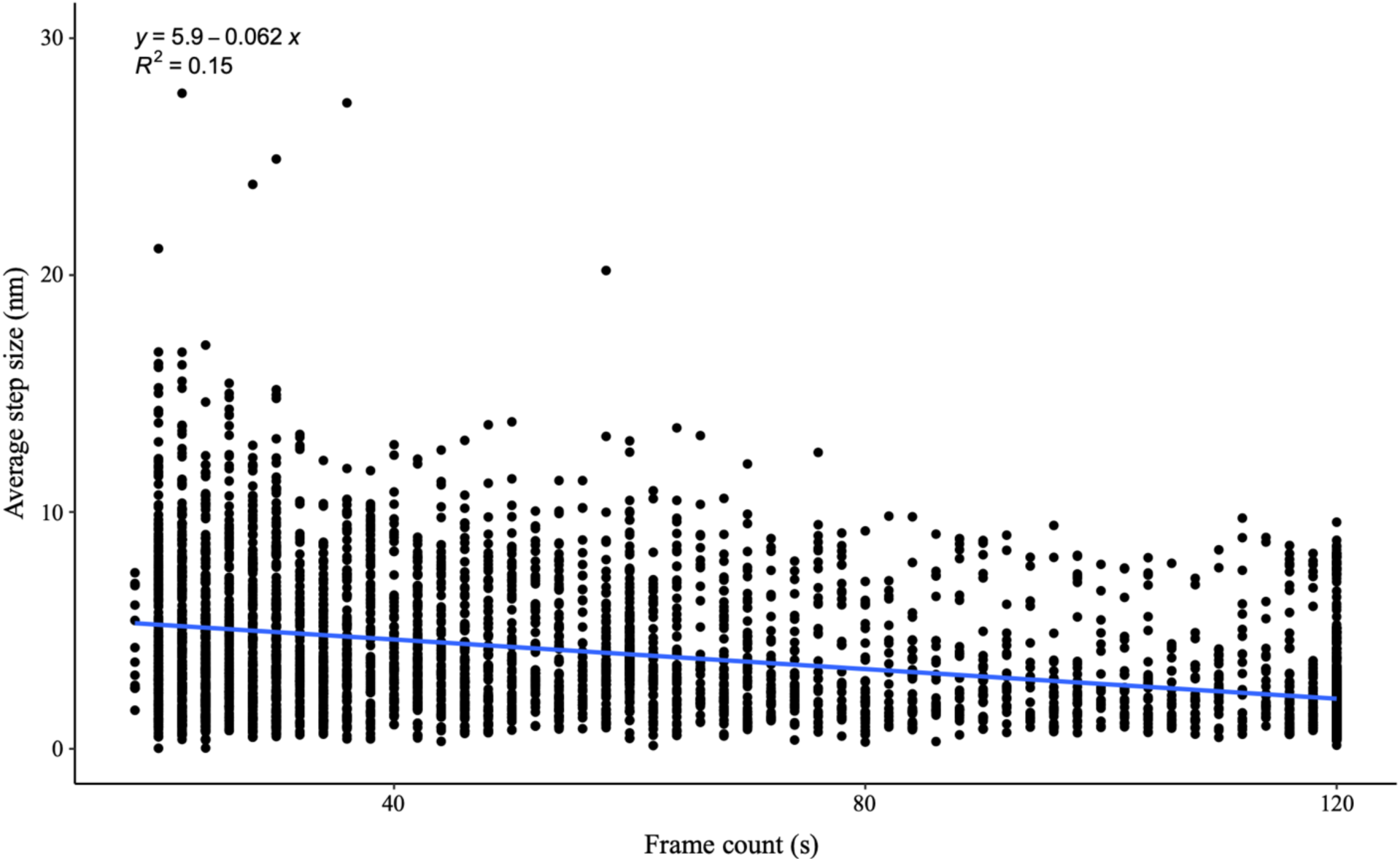
The scatter plot of average step size over each frame of the movies shows no bias by the AFM tip, as the R^2^ linear regression is 0.15. The data is obtained from two independent technical replicates per condition.

**Supplemental Figure 5.**
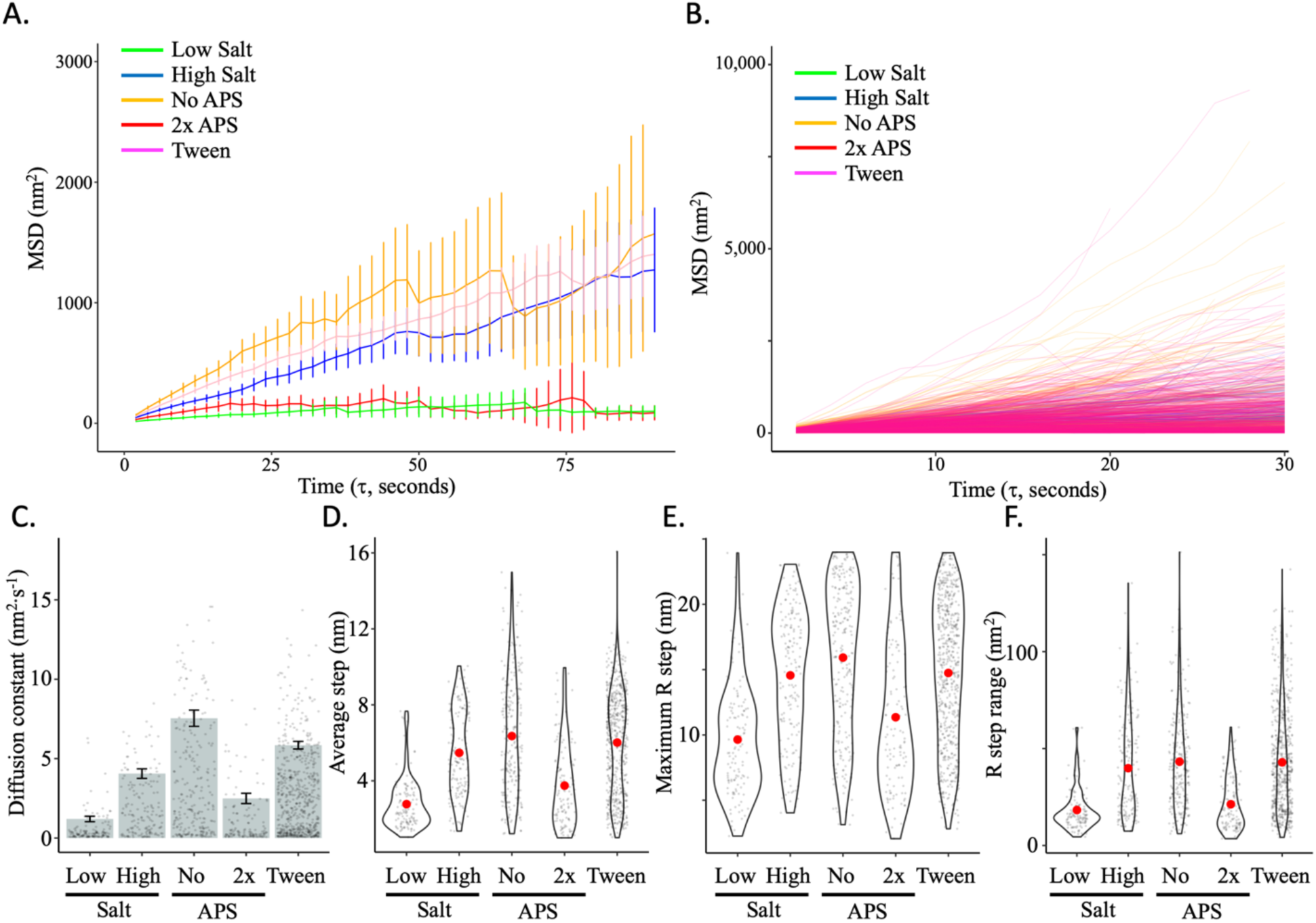
(A) The average mean square displacement (MSD) is shown with standard error as a function of the time interval for all control conditions (low salt = green; high salt = blue; no APS = yellow; 2x APS = red; 0.01% Tween = pink). (B) Individual MSD curves for all control conditions. (C) The diffusion constants obtained from the MSD curves, with an individual data point for each trajectory shown. (D) The average single frame step-size for each individual tracked nucleosome (grey points). (E) The maximum R step for each individually tracked nucleosome per control condition. (F) The R step range for each individually tracked nucleosome per control condition. Red dots represent the overall mean. The line and bar graphs represent two independent technical replicates per condition. The error bars represent standard error of the mean.

**Supplemental Figure 6.**
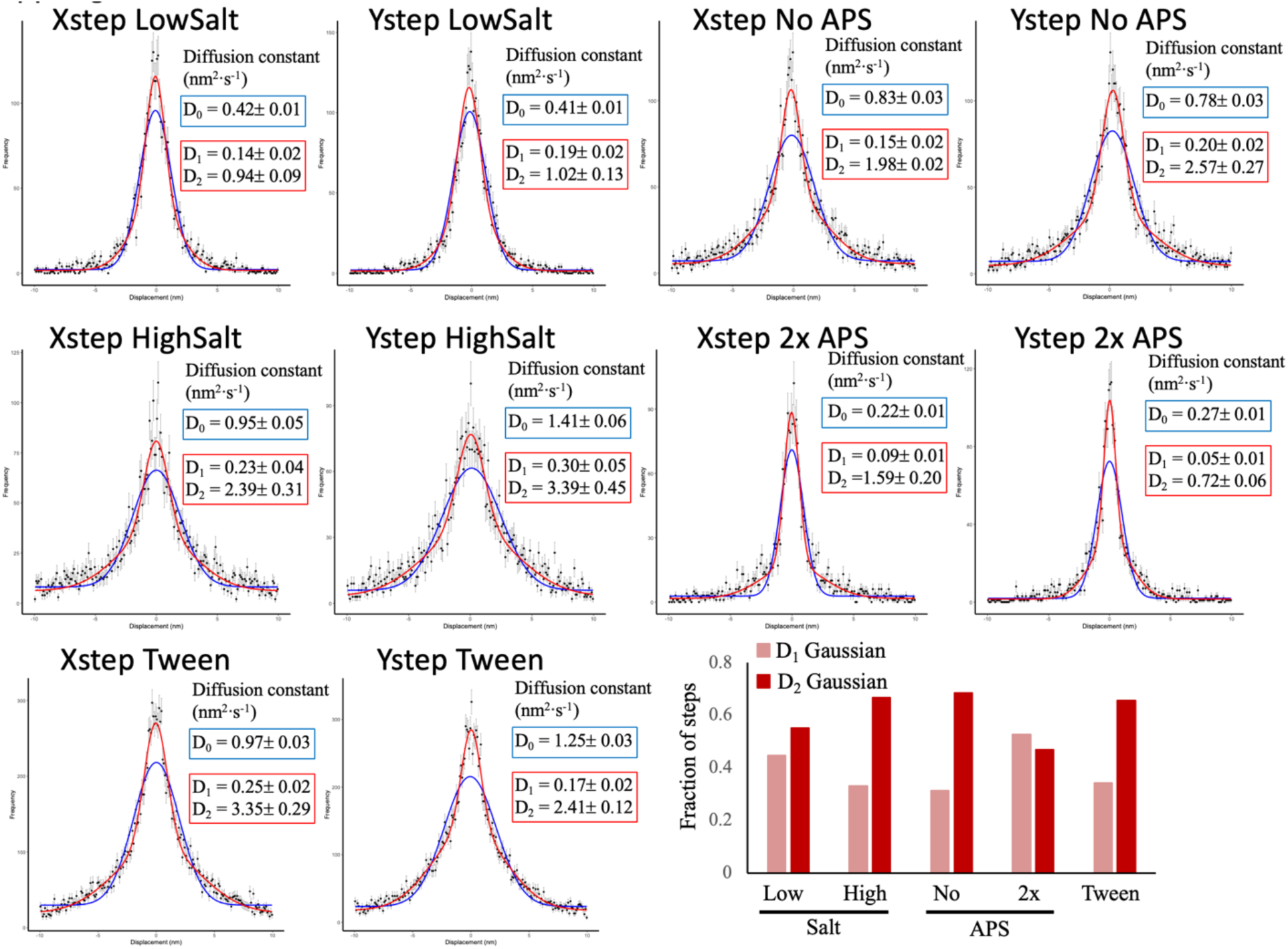
Single and double (Sum of two) Gaussian fitting of the x and y step displacement distributions for each of the control conditions. The double Gaussian distribution best fitted the x and y step displacement for all samples. The relative fraction of the D1 and D2 Gaussian fit are shown, where the D2 (larger standard deviation, higher diffusion constant) Gaussian is most prevalent for all conditions except 2x APS. The data is obtained from two independent technical replicates per condition.

**Supplemental Figure 7.**
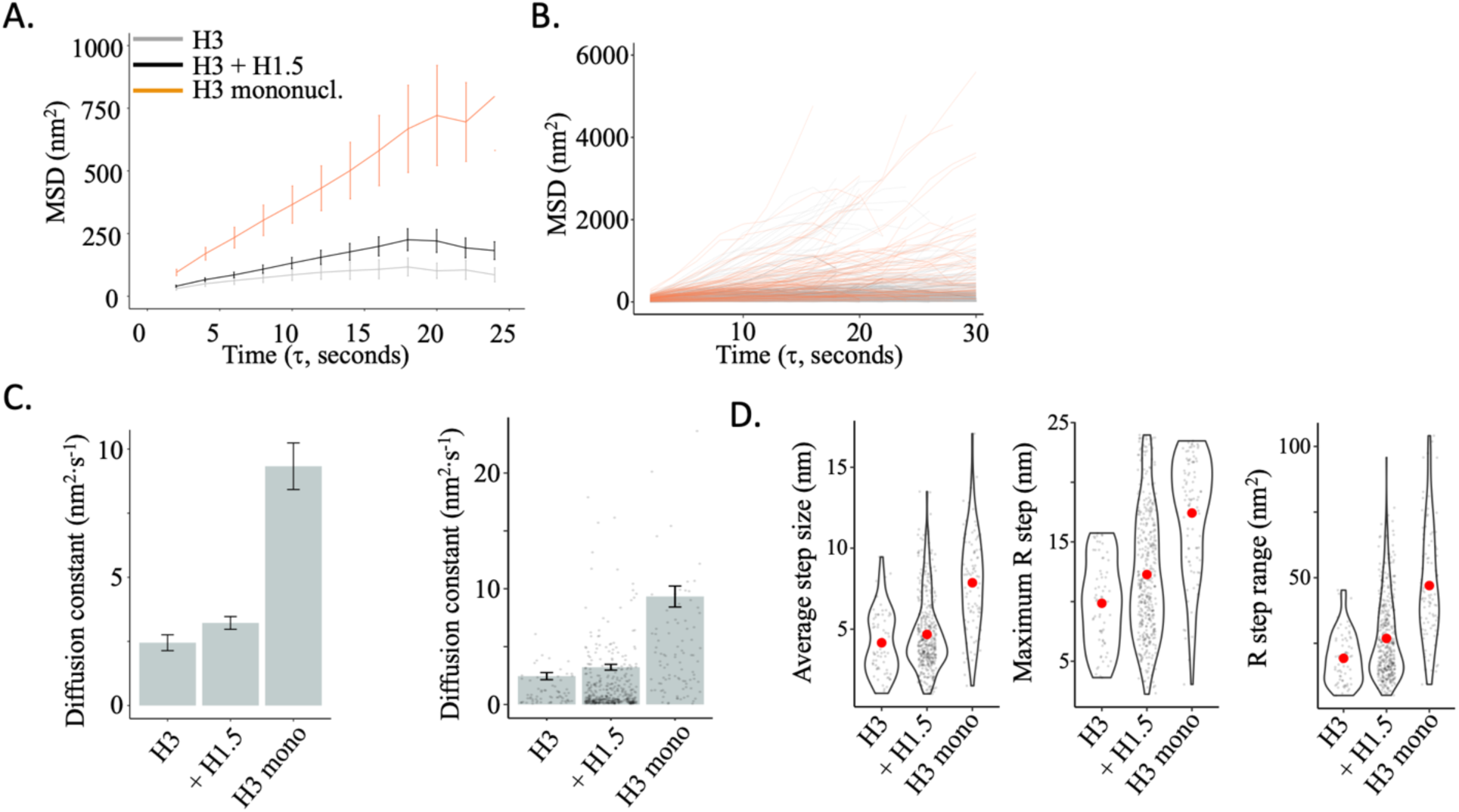
(A) The average (MSD) is shown with standard error as a function of the time interval for H3 chromatin (grey) alone or with linker histone H1.5 (black) (movies from Melters & Dalal 2020 JMB) and H3 mononucleosomes (orange). (B) Individual MSD curves for the three H3 samples. (C) Diffusion constants obtained from the MSD curves, with individual data point for each trajectory shown. The error bars represent standard error of the mean. (D) The average single frame step-size for each individual tracked nucleosome. The maximum R step for each individually tracked nucleosome. The R step range for each individually tracked nucleosome. Red dots represent the overall mean. The line and bar graphs represent at least two independent technical replicates.

**Supplemental Figure 8.**
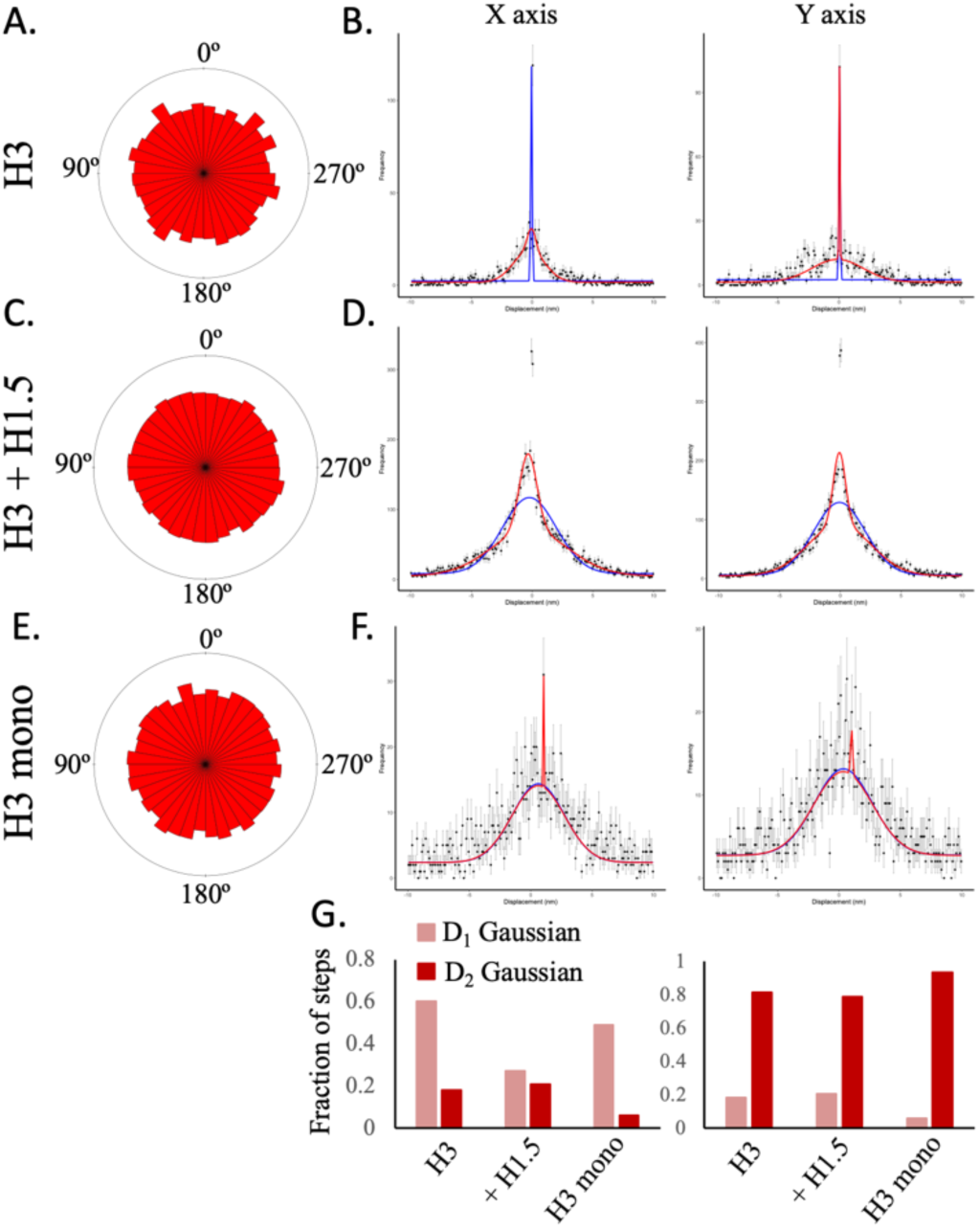
(A) The distribution of the angle between successive steps for H3 chromatin. (B) The single (blue line) and double (red line) Gaussian fitting of the x and y step displacements for H3 chromatin are shown. The double Gaussian distribution provided the best fit to the x step displacement but not the y step displacement distributions. The sharp peak represents stuck or immobile particles. (C) The distribution of the angle between successive steps for H3 chromatin. (D) The single and double Gaussian fitting of the x and y step displacement for H3 chromatin with H1.5 (added at 0.2 ratio H1 to H3 nucleosome) are shown. The double Gaussian distribution best fitted the x and y step displacement distributions. (E) The distribution of the angle between successive steps for H3 chromatin. (F) The single and double Gaussian fitting of the x and y step displacement for H3 mononucleosomes are shown. The single Gaussian performed as well as the double Gaussian fitting. The sharp peak off center is indicative of an artifact. (G) Ratio of D1 and D2 Gaussian fitting for all three samples. The data represent at least two independent technical replicates.

**Supplemental Figure 9.**
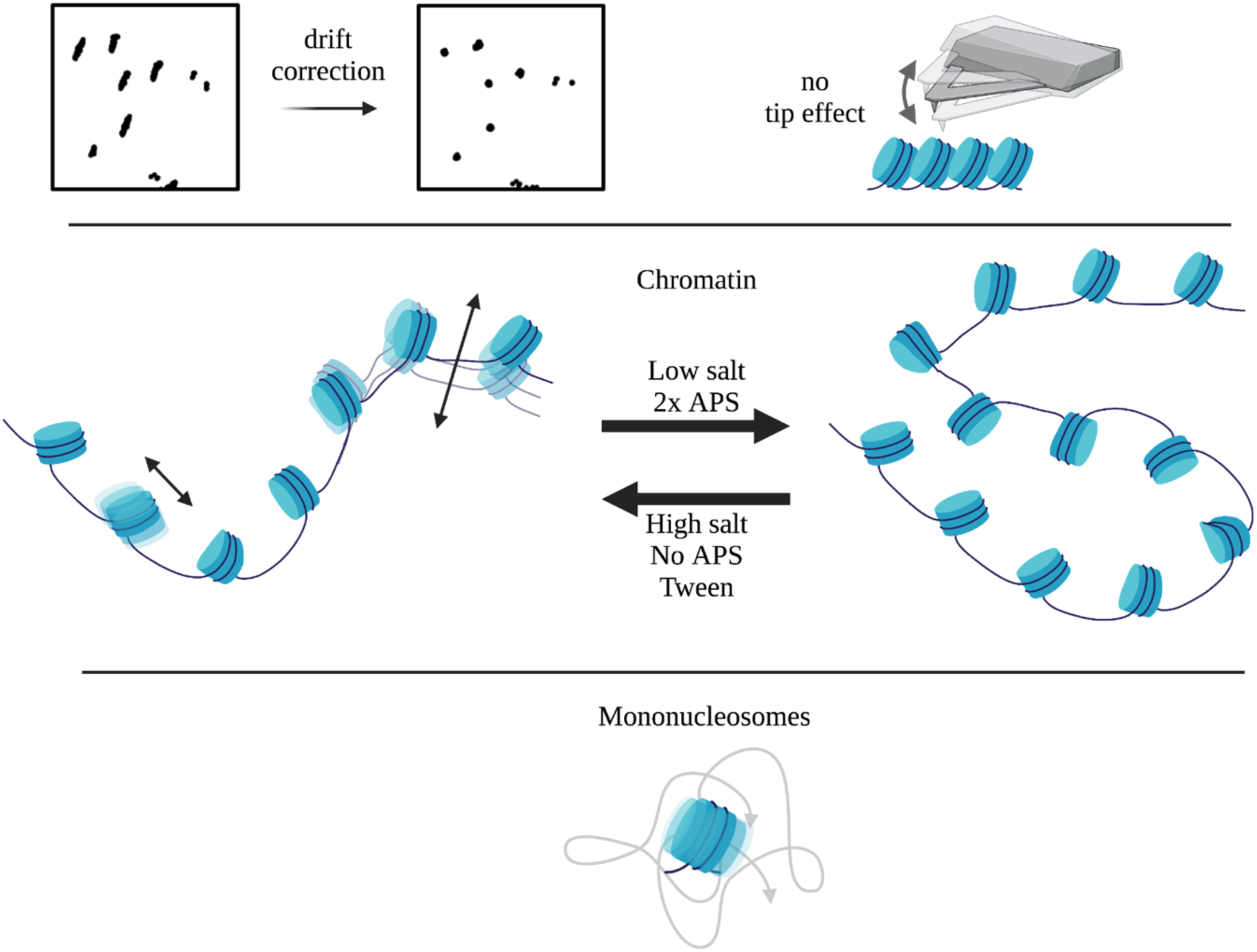
Schematic summary of the controls for HS-AFM quantitative single particle analysis. Movies were correct for drift. No tip-induced artifacts were observed. Low salt and 2x APS resulted in less mobile chromatin, whereas high salt, no APS, and Tween-20 conditions resulted in more mobile chromatin. H3 mononucleosomes were mobile without the constrains of being part of a polymer, resulting in similar single and double Gaussian fitting of single step displacement distributions.

**Supplemental Figure 10.**
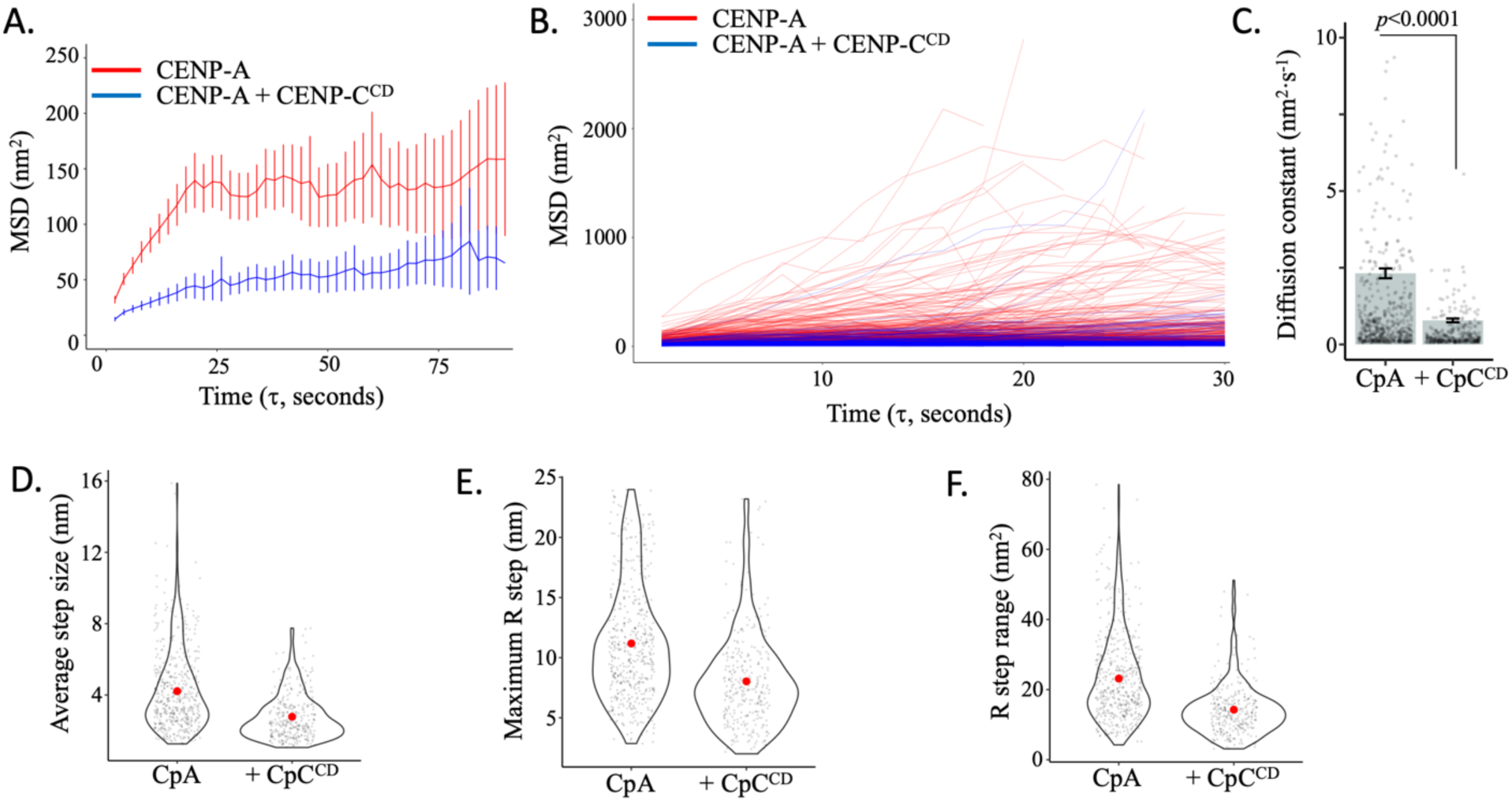
(A) The average mean square displacement (MSD) is shown with standard error as a function of the time interval for all CENP-A chromatin (red) and CENP-A chromatin with CENP-C^CD^ (blue, 2.2 molar excess to CENP-A nucleosomes). (B) Individual MSD for both samples. (C) The diffusion constants obtained from the MSD curves, with individual data points for each trajectory shown as grey points. Error bars represent standard error of the mean. (D) The average single frame step-size for each individual tracked nucleosome (grey points). (E) The maximum R step for each individually tracked nucleosome. (F) The R step range for each individually tracked nucleosome. Red dots represent the overall mean. The line and bar graphs represent at least three independent technical replicates.

**Supplemental Figure 11.**
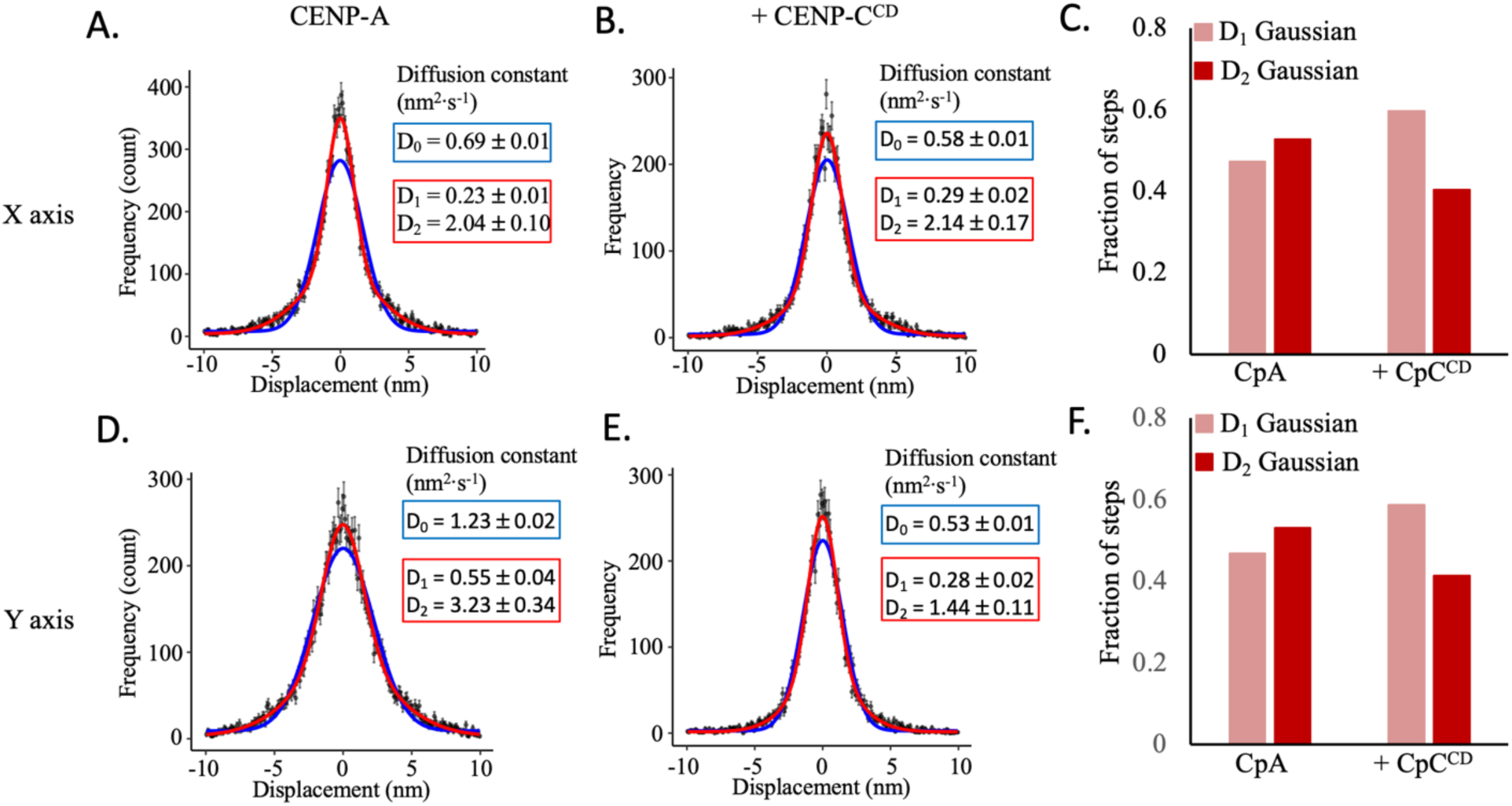
(A) The single and double Gaussian fitting of the x step displacement for CENP-A chromatin is shown. (B) The single and double Gaussian fitting of the x step displacement for CENP-A chromatin + CENP-C^CD^ is shown. (C) The double Gaussian distribution best fitted the x step displacement for both samples. The fraction of D1 and D2 Gaussians are shown, where the D2 Gaussian is most prevalent in most conditions except 2x APS. (D) The single and double Gaussian fitting of the y step displacement for CENP-A chromatin is shown. (E) The single and double Gaussian fitting of the y step displacement for CENP-A chromatin + CENP-C^CD^ is shown. (F) The double Gaussian distribution best fitted the y step displacement for both samples. The relative fraction of the D1 and D2 Gaussian fit are shown, where the D2 Gaussian is most prevalent in most conditions, except 2x APS. The data represent at least three independent technical replicates.

**Supplemental Figure 12.**
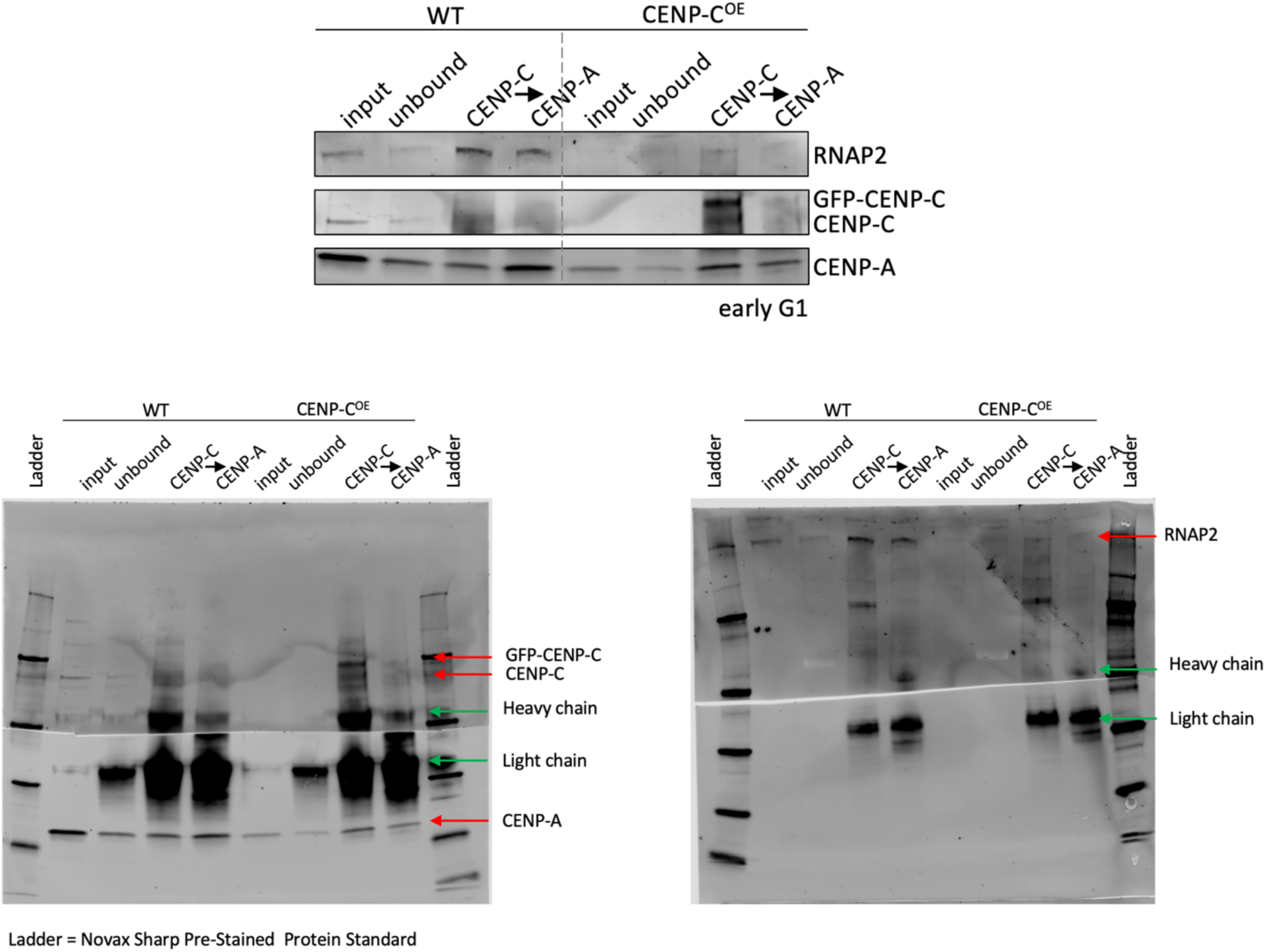
Representative western blot quantifying RNAP2 and CENP-A levels. HeLa cells were transfected with an empty vector (WT) or overexpressing CENP-C (CENP-C^OE^) and synchronized to early G1, when centromeric transcription is at its highest, similar to what we reported earlier (Melters *et al* 2019 PNAS). We probed for RNAP2, CENP-C, and CENP-A. The western blot is a representative blot of three independent technical replicates with representative original blots shown below.

**Supplemental movie 1** – HS-AFM movie of *in vitro* reconstituted CENP-A chromatin in 5 mM NaCl containing buffer (low salt; speed = 2x).

**Supplemental movie 2** – HS-AFM movie of *in vitro* reconstituted CENP-A chromatin in 150 mM NaCl containing buffer (high salt; speed = 2x).

**Supplemental movie 3** – HS-AFM movie of *in vitro* reconstituted CENP-A chromatin imaged without functionalized mica (no APS; speed = 2x).

**Supplemental movie 4** – HS-AFM movie of *in vitro* reconstituted CENP-A chromatin imaged with double the amount of APS to functionalize mica (2x APS; speed = 2x).

**Supplemental movie 5** – HS-AFM movie of *in vitro* reconstituted CENP-A chromatin imaged with physiological buffer + 0.01% Tween (Tween; speed = 2x).

**Supplemental movie 6** – HS-AFM movie of in vitro reconstituted H3 chromatin (speed = 2x).

**Supplemental movie 7** – HS-AFM movie of in vitro reconstituted H3 chromatin with H1.5, where H1.5 is added at a 0.2 molar ratio to H3 nucleosomes (speed = 2x).

**Supplemental movie 8** – HS-AFM movie of in vitro reconstituted H3 mononucleosomes (speed = 2x).

**Supplemental movie 9** – HS-AFM movie of *in vitro* reconstituted CENP-A chromatin (speed = 2x).

**Supplemental movie 10** – HS-AFM movie of in vitro reconstituted CENP-A chromatin with CENP-C^CD^ where CENP-C^CD^ is added at a 2.2x molar ratio to CENP-A nucleosomes (speed = 2x).

